# Cell type-specific enhancers regulate IL-22 expression in innate and adaptive lymphoid cells

**DOI:** 10.1101/2025.04.02.646834

**Authors:** Ankita Saini, Leone S. Hopkins, Vanida A. Serna, Matthew V. D. McCullen, Nicholas G. Selner, Bishan Bhattarai, José L. Fachi, Rebecca Glynn, Katharina E. Hayer, Craig H. Bassing, Marco Colonna, Eugene M. Oltz

## Abstract

IL-22, a signature cytokine of type 3 lymphoid cells, mediates epithelial homeostasis and protective pathogen responses in barrier tissues, while its deregulated expression drives chronic inflammation associated with colitis and psoriasis. Despite its therapeutic value, little is known about regulatory elements for IL-22 expression. We identify two conserved enhancers, E22-1 and E22-2, which differentially regulate *Il22* in type 3 lymphoid subsets. These enhancers are required for steady-state expression of gut antimicrobial peptides, protection from *C. rodentium* infection, and development of IL-22-mediated psoriasis. E22-1 resembles many known enhancers, functioning in both Th-ILC counterparts. However, E22-2 is only required for IL-22 expression in ILC3s. Its ILC3 restriction relies on multiple Runx3 sites, combined with the lack of a functional RORγt motif, which is present in E22-1. Thus, although responding to similar stimuli, type 3 lymphoid cells use distinct cis-elements for IL-22 expression, with E22-2 likely serving as a homeostatic enhancer in barrier tissues.

## Introduction

The cytokine IL-22 functions as an important regulator of epithelial homeostasis and antimicrobial responses in barrier tissues, including the gastrointestinal tract, lung, liver, skin, and breast^1–5^. The primary sources of IL-22 are type 3 innate lymphoid cells (ILC3s) and their adaptive counterparts, Th17/22 cells, which are collectively referred to as type 3 immune cells since they have similar effector programs^1,4,6–9^. In addition, IL-22 is produced by other innate cells, such as NKT17 and γδ-T cells in response to danger signals associated with infection or inflammation. The cognate receptor for IL-22 is a heterodimer of IL-22R1 and IL-10R2, which is expressed on epithelial and stromal cells, as well as other cells of non-hematopoietic origin^10^. Binding of IL-22 to its receptor primarily triggers the JAK1-STAT3 signaling pathway, which induces expression of anti-microbial peptides (AMPs), chemokines, anti-apoptotic molecules, and genes involved in cell proliferation, tissue repair, and regeneration^11^. As such, IL-22 is pivotal in the initiation of antimicrobial responses, resolution of inflammation, and restoration of barrier integrity.

In this regard, IL-22 and its sources play a multi-faceted role in maintaining intestinal health and defense against pathogens that infect gut tissues, processes that have been especially well-studied in mice infected by *Citrobacter rodentium*^8,12–15^. This enteric bacterium attaches to mucosal surfaces in the colon, disrupting barrier function, and it serves as a pathogenesis model for *E. coli*-induced colitis in humans. Indeed, *C. rodentium* infection in IL-22 deficient mice leads to severe colitis and high mortality rates^12^. Recent studies have shown that IL-22 derived from innate lymphoid cells, such as ILC3s, plays a crucial role in controlling the early stages of *C. rodentium* infection, while antigen-specific CD4^+^ Th17/22 populations expand and become important sources of IL-22 during later stages of infection, aiding in final clearance of the pathogen^13,14^.

Conversely, perturbations in IL-22 levels have been associated with several cancers, as well as inflammatory and autoimmune disorders, emphasizing the pathogenic role of IL-22^16,17^. In patients suffering from psoriasis, a chronic skin inflammatory disorder, enhanced levels of IL-22 in skin and blood have been correlated with disease severity^18^. Likewise, in mice, imiquimod (IMQ) induces a psoriasis-like skin inflammation that is driven by IL-22 and IL-17^19–21^. However, the relevant contribution of innate and adaptive immune cell types producing these pathogenic type 3 cytokines remains unresolved. In this regard, many studies have underscored the potential of IL-22 as a therapeutic target for chronic inflammatory conditions involving barrier tissues and ongoing clinical trials are focused on drugs that either impact the production, neutralization, or downstream signaling of IL-22^22^.

While the effects of IL-22 on epithelial cells have been well-studied, significant gaps remain in our understanding about cis-regulatory circuits that control homeostatic IL-22 expression or its induction in lymphoid cells during normal or pathogenic responses. At least some ILC3s express IL-22 at a steady state in healthy tissues, whereas CD4^+^ T helper cells normally require TCR and/or cytokine activation for its expression. IL-23 is the major cytokine that induces IL-22 in both innate and adaptive immune cells, which, in conjunction with IL-1β, leads to optimal IL-22 production^12,17,23,24^. These pathways activate STAT3 and the signature type 3 transcription factor, RORγt, which are known to augment the expression of *Il22* at the RNA level. However, understanding how cis-regulatory circuits coordinate disparate signals to tune *Il22* gene expression in different type 3 immune cells inhabiting normal and pathogenic microenvironments remains a key goal, especially given the therapeutic potential of regulating *Il22* expression.

To address this knowledge gap, we performed a CRISPRi screen of potential cis-regulatory elements embedded within the 1.5 Mb *Mdm1*-*Il22*-*Ifng* locus using an ILC3 cell model. We identified two conserved enhancers, E22-1 and E22-2, which differentially regulate *Il22* in innate versus adaptive lymphoid cells. Together, these enhancers are required for homeostatic, protective, and pathogenic IL-22 expression in vivo. Remarkably, E22-2 element is highly specific for ILC3s and a small subset of γδ-T cells. This activity profile distinguishes E22-2 from other known type 3 enhancers, which normally function in both Th and ILC counterparts. Mechanistic studies revealed the underlying architecture of E22-2 that restricts its activity to ILC3s, likely to support its service as a homeostatic enhancer of IL-22 expression in barrier tissues.

## Results

### CRISPRi screen identifies potential cis-elements for *Il22* expression in type 3 lymphoid cells

In both humans and mice, *IL22* is situated in a locus that also contains genes encoding the type 1 cytokine IFN-γ, IL-26 (only in humans), and the cell cycle regulator *MDM1* **(Fig. 1a)**. These multigenic loci are characterized by a complex regulatory space with many conserved DNA elements that have active chromatin characteristics in lymphoid cells, including a large ILC3-specific super-enhancer (SE) region in both species (**Fig. 1b**)^7^. To determine which of these accessible chromatin regions are necessary for regulating *Il22* expression in type 3 cells, we performed a CRISPRi screen in the mouse ILC3 cell model, Mnk3^25^. Like primary ILC3s, Mnk3 cells produce high levels of IL-22 and IL-17 in response to the type 3 agonists IL-1β and IL-23. Moreover, accessible chromatin patterns in Mnk3, as observed in new ATAC-seq data, largely paralleled those reported previously for primary mouse ILC3s (**Fig. 1b**)^26,27^.

**Fig. 1:**
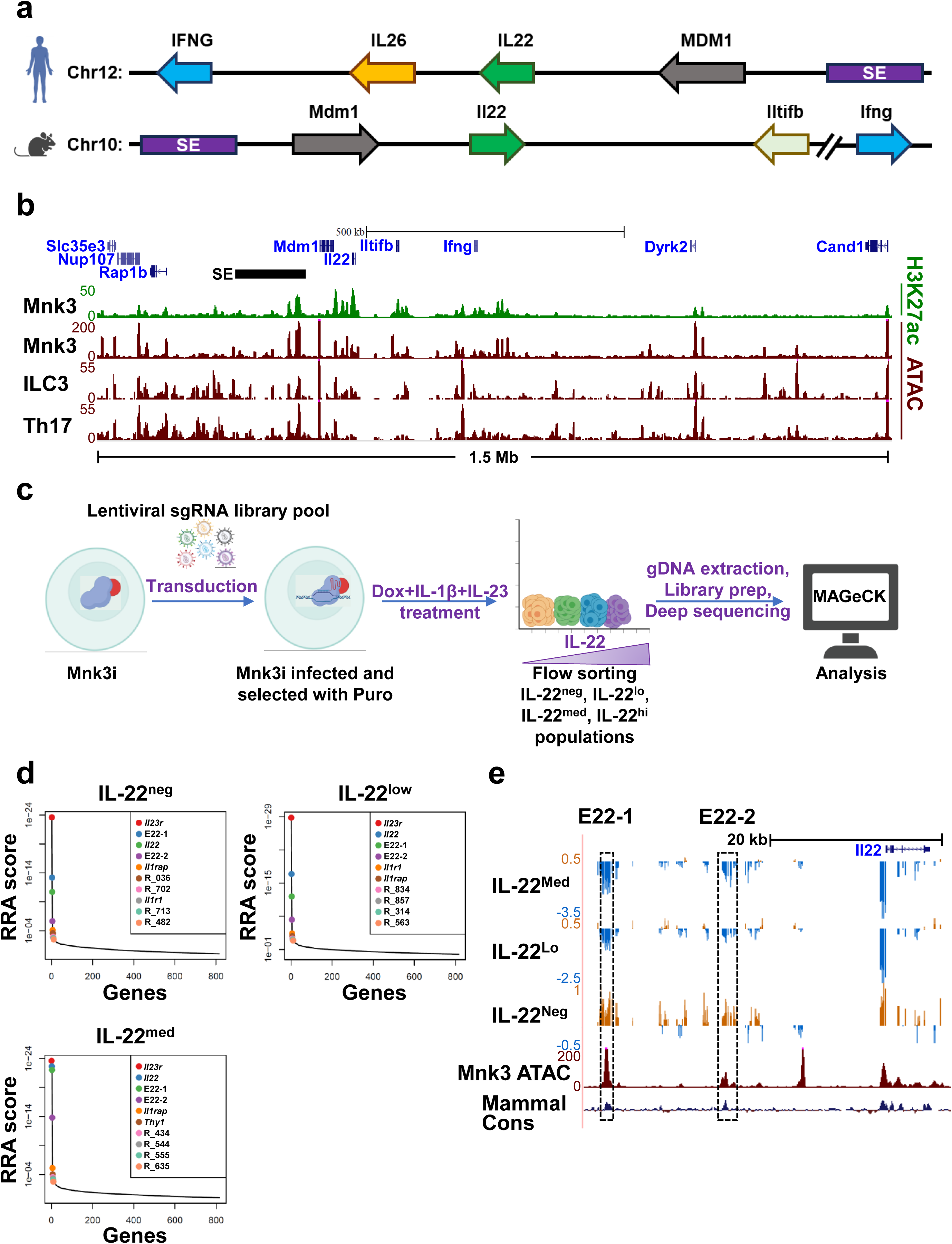
CRISPRi screen identifies potential *Il22* enhancers in Mnk3 cells. (a) IL-22 gene locus in human (top) and mouse (bottom). SE designates a lymphoid-specific super-enhancer. (b) UCSC genome browser tracks (mouse, mm9) for ATAC-(maroon) or H3K27ac-seq (green) data in Mnk3 cells stimulated with IL-1β+IL-23, primary ILC3s, and Th17 cells. (c) Schematic of CRISPRi screen design. (d) MAGeCK’s Robust Rank Aggreation (RRA) distribution plots in +dox cellular populations showing top 10 regions with sgRNA enrichment in IL-22 negative or with depletion in IL-22 low or medium cell fractions when compared with the corresponding -dox controls. Targets designated R_xxx are regions within the *Il22* locus which were not enriched/depleted in all three sorted fractions (Table S3). (e) UCSC genome browser view (mm9) of the *Il22* locus showing log fold change in sgRNA depletion (blue) or enrichment (gold) in the indicated populations sorted for IL-22 expression levels when treated with dox (dCas9-KRAB^hi^) versus their corresponding –dox counterparts.

To perform CRISPRi screens, we engineered a Mnk3 subclone that expresses tetracycline inducible catalytically inactive Cas9 associated with a KRAB chromatin repressor domain (dCas9-KRAB) and an mCherry reporter (Mnk3i cell line). The Mnk3i line was infected with a custom lentiviral sgRNA library (MOI < 0.3) that tiles all accessible chromatin regions in a ∼1.5 Mb space surrounding the *Il22* locus (**Fig. 1c**). The accessible regions chosen for this library were identified using ATAC-seq data from a collection of different lymphoid cell types, including Mnk3, the ILC1 model Mnk1, primary type 1 cells (NK, ILC1, and Th1), and primary type 3 cells (ILC3 and Th17)^26,27^. The lentiviral library also had a number of control sgRNAs that targeted the *Il22, Il23r,* and *Il1rap* promoters (positive controls), as well as ATAC-negative regions on chromosome 10 and sgRNAs with no targets in the mouse genome (negative controls). Lentiviral-infected cells were selected with puromycin, followed by treatment with doxycycline (dox) (24 hours), before adding the type 3 agonists IL-1β and IL-23. After 16 additional hours, cells were flow sorted into populations that expressed high, medium, low, or no detectable IL-22. sgRNA sequences were amplified by PCR from each fraction, which were deep sequenced and analyzed by the MAGeCK pipeline^28^. The fraction expressing the highest levels of IL-22 yielded low cell numbers and was excluded from further analysis.

Because this was an inhibitory screen, we focused on regions in which sgRNAs were enriched significantly in the IL-22 negative fraction but were depleted in cells expressing IL-22, when compared with sgRNA representation in their counterpart fractions that were not treated with dox (i.e., lacking dCas9-KRAB). The top 10 regions exhibiting these characteristics are shown in **Fig. 1d**. As expected, sgRNAs targeting promoters of *Il22*, *Il23r*, and *Il1rap*, a component of IL-1β receptor complex, were in this top 10 list, whereas sgRNAs targeting ATAC-negative regions were neither significantly enriched nor depleted in any cell population (**Fig. 1d,e and S1a-c**). Among the top 10 hits, only two regions showed the expected enrichment-depletion profiles in all three sorted cell fractions, as well as high sequence conservation in mammals. We named these accessible chromatin regions, E22-1, positioned ∼32 kb upstream, and E22-2, located ∼17 kb upstream of the *Il22* promoter (**Fig. 1e**). Of note, no accessible regions within the ILC3-specific SE were identified using this CRISPRi approach, potentially because of functional redundancy within the SE, or because the SE may be irrelevant for IL-22 expression in this cell model. Notwithstanding, the CRISPRi screen identified E22-1 and E22-2 as candidate cis-elements that regulate *Il22* expression in type 3 immune cells.

### E22-1 and E22-2 function as *Il22* enhancers in Mnk3 cells

To further explore whether E22-1 and E22-2 serve as *Il22* regulatory elements, we surveyed chromatin profiles in several relevant lymphoid populations (Mnk3, NK, ILC1, ILC3, and Th17 cells), using new or published ATAC- and H3K27ac-seq data^26^. The latter histone modification represents an excellent metric for enhancer activity. As shown in **Fig. 2a**, the distal element, E22-1, was highly accessible in type 3 (ILC3, Th17, and Mnk3) but not in type 1 cells (NK and ILC1), and stimulation of Mnk3 with IL-1β+IL-23 further enhanced its H3K27ac. By contrast, E22-2 was accessible only in ILC3s, but not in Th17 or type 1 cells. Like E22-1, H3K27ac at E22-2 was enhanced in Mnk3 upon IL-1β+IL-23 treatment. These chromatin data suggest that both E22-1 and E22-2 are enhancers that can be induced by type 3 agonists.

**Fig. 2:**
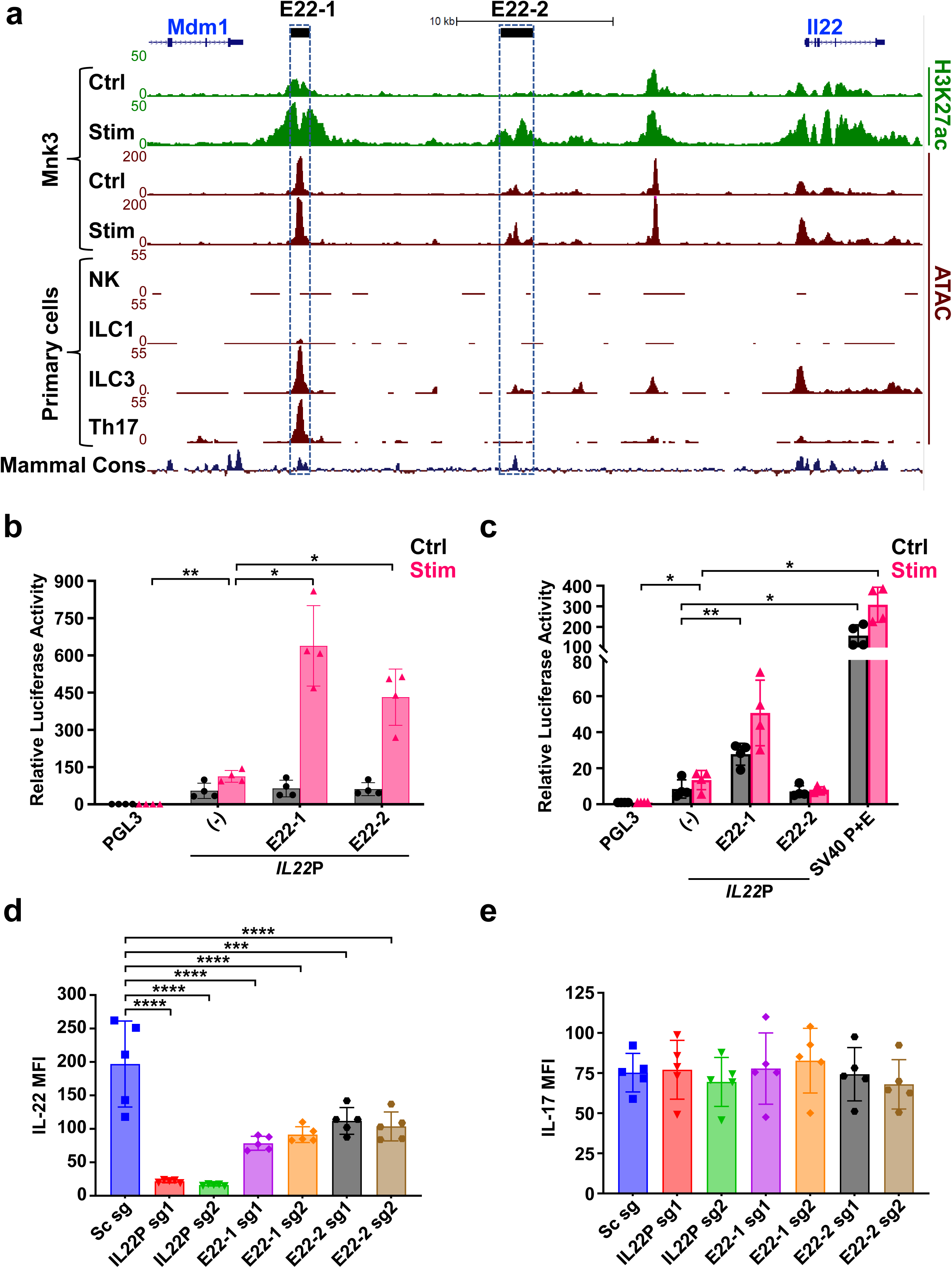
E22-1 and E22-2 regulate *Il22* expression in Mnk3. (a) UCSC genome browser view of ATAC-seq (maroon) or H3K27ac-seq (green) data for Mnk3 cells (untreated or treated with IL-1β+IL-23), and for primary mouse cells (NK, ILC1, ILC3, and Th17). A mammalian conservation track is shown at the bottom. E22-1 and E22-2 are boxed. (b) Luciferase reporter assays for enhancer activity of E22-1 and E22-2 in Mnk3 cell after nucleofection with the indicated vectors (PGL3 backbone, *Il22* promoter alone (-), or in combination with each enhancer). After 24 h, cells were either treated with 10 ng/ml each of IL-1β and IL-23 or left untreated for 16 h and harvested to assess reporter activity. (c) Nucleofected EL4 cells were rested overnight, then either treated with 1X cell stimulation cocktail or left untreated for 4 h and harvested. Reporter activities relative to PGL3-Basic vector are shown, in addition to results from a positive control vector containing the SV40 viral promoter/enhancer (PGL3-Control Vector). (d, e) Flow cytometric analysis showing mean fluorescence intensity (MFI) of (d) IL-22 and (e) IL-17F. The Mnk3i line was transduced with viral particles containing sgRNAs targeting the *Il22* promoter, E22-1, E22-2, or a scrambled sgRNA, followed by puromycin selection. Transduced cells were treated with dox for 24 h, followed by overnight stimulation with 10 ng/ml each of IL-1β and IL-23, and stained intracellularly for IL-22 and IL-17F. Flow cytometry and luciferase data are shown as mean ± SD and are pooled from (b, c) four independent experiments or (d, e) five independent experiments. \**P* < 0.05; \*\**P* < 0.01; \*\*\**P* < 0.001; \*\*\*\**P* < 0.0001; two-way ANOVA with Dunnett’s multiple comparisons test (b, c) or one-way ANOVA with Dunnett’s multiple comparisons test (d, e).

To test for enhancer function, we performed luciferase reporter assays on cloned E22-1 and E22-2 sequences in Mnk3 cells using vectors driven by the mouse *Il22* promoter. Neither element augmented expression from the *Il22* promoter in unstimulated Mnk3 cells, but treatment with IL-1β+IL-23 induced robust enhancer activity for both E22-1 (∼4-fold) and E22-2 (∼3-fold) (**Fig. 2b**). Consistent with chromatin profiling, only E22-1 displayed enhancer activity in the mouse T-cell line, EL4 (**Fig. 2c**). As expected, neither element enhanced reporter activity in the mouse B-cell line, 63-12 (**Fig. S2a**). To further establish their enhancer functions at the endogenous *Il22* locus, we introduced sgRNAs that were designed to target either E22-1 or E22-2 in Mnk3i. Induction of dCas9-KRAB mediated significant repression of IL-22 with either enhancer sgRNA compared with a scrambled sgRNA in stimulated Mnk3 cells (**Fig. 2d**), but none of the sgRNAs affected levels of the other type 3 cytokine, IL-17F (**Fig. 2e**), whose gene is located on a different chromosome. In addition, expression of the neighboring gene, *Mdm1*, was unaffected by CRISPRi-mediated repression of E22-1 or E22-2, whereas *Il22* transcripts were significantly diminished (**Fig. S2b,c**).

In some mouse strains, including C57BL/6, 129 and FVB, the region between *Il22* and *Ifng* harbors a pseudogene ∼80 kb downstream of *Il22*, called *Iltifb*, which is an inverted duplication of *Il22^29^*. The *Iltifb* region also contains a duplicate of E22-2, which we coined twin-E22-2, but has no duplicate of E22-1 (**Fig. S2d**). Our CRISPRi screen did not identify twin-E22-2 as a putative candidate regulating *Il22*. Nevertheless, the similarities between E22-2 and twin-E22-2 merited direct verification to rule out any role for the latter in *Il22* gene regulation. For this purpose, we designed sgRNAs that specifically target either E22-2 or twin-E22-2 and verified their specificities using in vitro cutting assays with Cas9 ribonucleoprotein complexes (**Fig. S2e**). IL-22 expression was unchanged in a Mnk3i line that stably expressed the twin-E22-2 sgRNA (**Fig. S2f)**. In contrast, Mnk3i cells expressing the highly specific E22-2 sgRNA exhibited a significant reduction in IL-22 levels. As expected, none of the sgRNAs impacted IL-17F expression (**Fig. S2g**). Collectively, our molecular assays in Mnk3 cells implicate E22-1 and E22-2 as the two most important cis-regulatory elements for *Il22* expression in this type 3 lymphoid cell model.

### In vivo regulation of *Il22* by E22-1 and E22-2

To investigate their functions in vivo, we generated mice lacking either E22-1(E22-1^Δ^) or E22-2 (E22-2^Δ^) or lacking both elements (E22-1/2^Δ^) using CRISPR-Cas9 technology. The E22-1^Δ^ and E22-2^Δ^ mice were created by deletion of 3.5 and 2.5 kb, respectively, spanning regions covered by H3K27ac in Mnk3 cells. The E22-1/2^Δ^ strain was generated by deleting E22-1 from E22-2^Δ^ mice using the same sgRNAs. All deletions were confirmed by DNA sequencing and the double knockout was verified to be on the same allele using long-range PCR.

To verify that E22-1 and E22-2 were primary IL-22 enhancers for type 3 cells in vivo, we first focused on mice lacking both elements. ILC3s were analyzed from small intestine lamina propria (SI-LP), which contains all three known subsets, namely, NKp46^+^, CCR6^+^ (lymphoid tissue inducer cells, LTi), and double negative (DN)^30^. We observed no significant changes in the frequency of ILC3 subsets in the E22-1/2^Δ^ mice compared to WT controls (**Fig. S3a**). Bulk SI-LP cells were cultured in the presence or absence of IL-1β and IL-23 for 3.5 h and IL-22 levels were quantified in each ILC3 subset using flow cytometry. As shown in **Fig. 3a-c**, mice lacking both elements had a ∼3-fold reduction in average IL-22 expression, as well as a significant reduction in the percentages of IL-22 positive cells for all three ILC3 subsets (**Fig. S3b-d**). To test how deletion of the two dominant enhancers affected IL-22 expression by Th22 cells, we cultured splenic naïve CD4^+^ T cells under Th22 polarizing conditions for 3 days. As shown in **Fig. 3d**, there was a dramatic reduction in both levels of IL-22 expression (∼4-fold) and percentages of IL-22 positive cells (∼7-fold; **Fig. S3f**) for Th22 cells derived from E22-1/2^Δ^ mice compared with WT animals. Importantly, IL-17 expression remained unaffected (**Fig. S3e**).

**Fig. 3:**
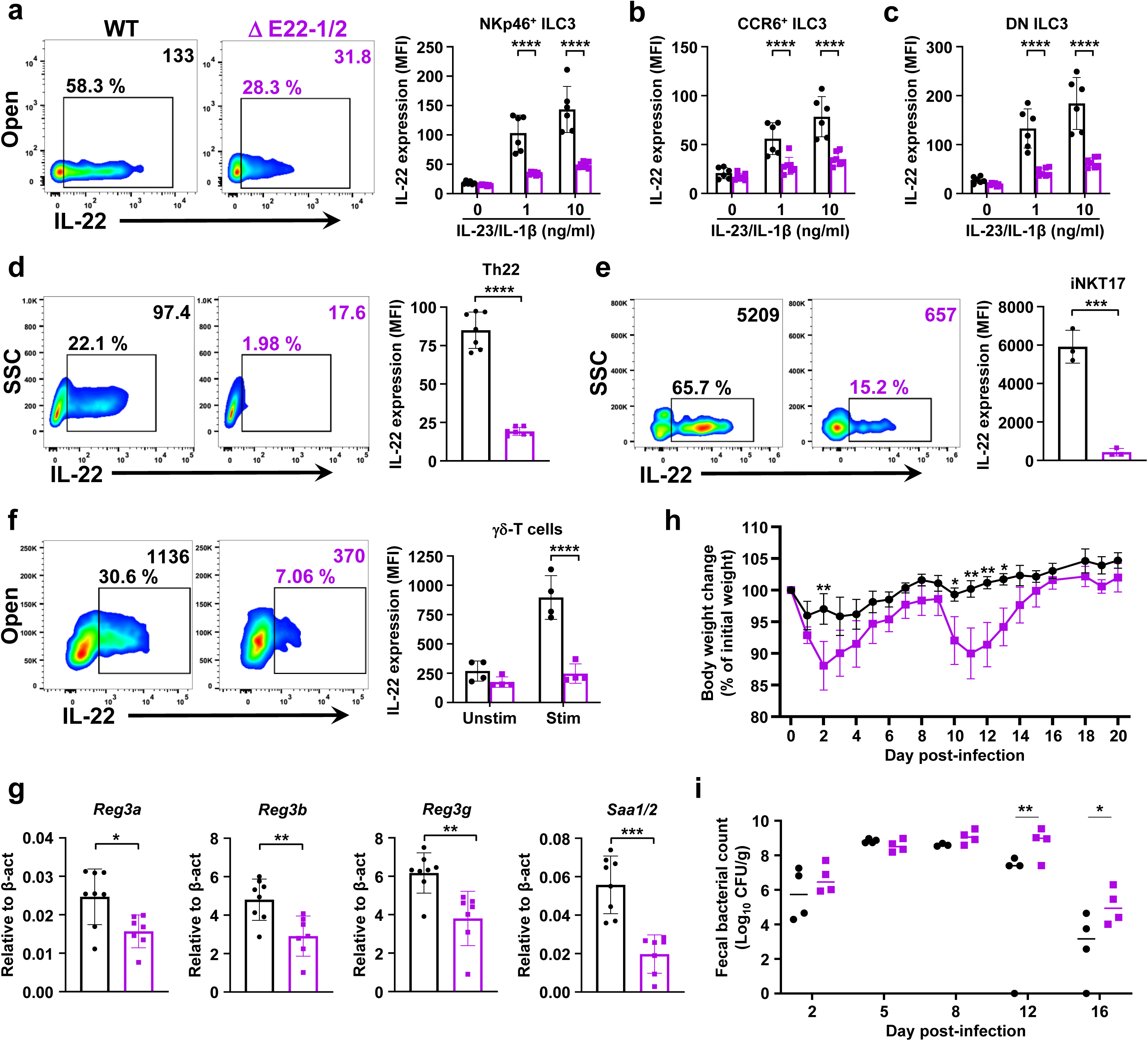
E22-1 and E22-2 are necessary for IL-22 production and type 3 immune functions in mice. (a-c) Representative flow plots (left) and quantification (right) of IL-22 MFI in SI-LP NKp46^+^ (a), CCR6^+^ (b), and DN ILC3 (c) subsets of WT (black) and E22-1/2^Δ^ mice (purple), after in vitro stimulation with indicated concentrations of IL-23+IL-1β for 3.5 h. (d-f) Representative flow plots (left) and expression of IL-22 (MFI) in ex vivo polarized Th22 (d), lymph node iNKT17 (e), and SI-LP γδ-T cells (f). (g) RT-qPCR analysis of IL-22 controlled AMP genes *Reg3a*, *Reg3b*, *Reg3g*, and *Saa1/2* in terminal ileum of WT and E22-1/2^Δ^ mice. Expression relative to *β-actin*. (h, i) Weight loss (h) and bug burden (CFU) in feces (i) of WT and E22-1/2^Δ^ mice infected with *C. rodentium* (2x10^9^ CFUs per animal). Data are shown as mean ± SD (a-g), mean ± SEM (h), or median (i) and are pooled (a-d, f, g), or are representative (e, h, i) of at least two independent experiments. \**P* < 0.05; \*\**P* < 0.01; \*\*\**P* < 0.001; \*\*\*\**P* < 0.0001; two-way ANOVA with Sidak’s multiple comparisons test (a-c, f, h), unpaired t-test (d, e, g), or two-way ANOVA followed by Fisher’s test (i).

In addition to ILC3 and Th17/22 cells, IL-22 is produced by NKT17 and γδ-T cells, two other innate immune cell types^1,4^. To determine the impact of the E22-1/2^Δ^ mutation on the former subset, we isolated peripheral lymph nodes cells and subjected them to overnight stimulation with anti-CD3, anti-CD28 along with IL-1β and IL-23. Cultured cells were then analyzed by flow cytometry to establish IL-22 expression levels in NKT17 cells. As shown in **Fig. 3e** and **S3g,h** NKT17 cells from E22-1/2^Δ^ mice had more than 10-fold reduction in IL-22, while IL-17 expression was unaffected. Lastly, γδ-T cells from SI-LP isolates were analyzed after stimulation with IL-1β+IL-23. The E22-1/2^Δ^ cells had ∼4-fold reduction in IL-22 compared with γδ-T cells from WT controls (**Fig. 3f and S3i**).

To test whether the observed reductions in IL-22 expression by type 3 cells was physiologically relevant, we examined the impact for the dual enhancer deletion on IL-22 dependent homeostatic and defense programs in the gut. With regards to the former, IL-22 regulates steady-state expression of AMPs by intestinal epithelial cells. As such, we performed RT-qPCR analysis for transcripts corresponding to several AMPs using RNA isolated from terminal ileum of WT and E22-1/2^Δ^ mice. We observed significantly reduced steady-state levels of transcripts corresponding to IL-22 inducible AMP genes, *Reg3*α, *Reg3*β, *Reg3*γ, and *Saa1/2*, in E22-1/2^Δ^ mice (**Fig 3g**). To test the functional importance of the E22-1 and E22-2 enhancers in mucosal defense, we infected WT and E22-1/2^Δ^ mice with *C*. *rodentium*. Over the course of 20 days following infection, E22-1/2^Δ^ mice experienced more pronounced weight loss at early (days 2-4) and late time points (days 10-14) as compared to WT controls (**Fig. 3h**). The weight loss was also accompanied by a significantly higher fecal bacterial burden on days 12 and 16 post-infection (**Fig. 3i**). Thus, the E22-1 and E22-2 enhancers are required for attaining levels of IL-22 expression that mediate both steady-state homeostatic protection in the gut and defense against a mucosal pathogen, *C. rodentium*.

### E22-1 and E22-2 both contribute to cell type-specific patterns of IL-22 expression and protection from imiquimod-induced psoriasis

To investigate the individual roles of E22-1 and E22-2, we first evaluated IL-22 production by type 3 immune cells in single knockout mice. We did not observe any significant change in the frequency of ILC3 subsets in either of the tested mutant mice (**Fig. S4a and l**). Deletion of E22-1 impaired expression of IL-22 in all three subsets of ILC3s, a reduction comparable to that observed in E22-1/2^Δ^ derived ILC3s (**Fig. 4a and S4b-f**). We also observed a significant decrease in the levels of IL-22 and the percentage of IL-22 positive Th22 cells after polarization of splenic naïve CD4^+^ T cells from E22-1^Δ^ mice, while IL-17 levels remained unchanged (**Fig. 4b and S4g,h**). In addition, both NKT17 and γδ-T cells from E22-1^Δ^ mice expressed reduced levels of IL-22 as compared to WT controls (**Fig. 4c,d and S4i-k**). Overall, the E22-1 mutation phenocopied the double knockout with regard to IL-22 expression in all tested type 3 lymphoid cell subsets.

**Figure 4:**
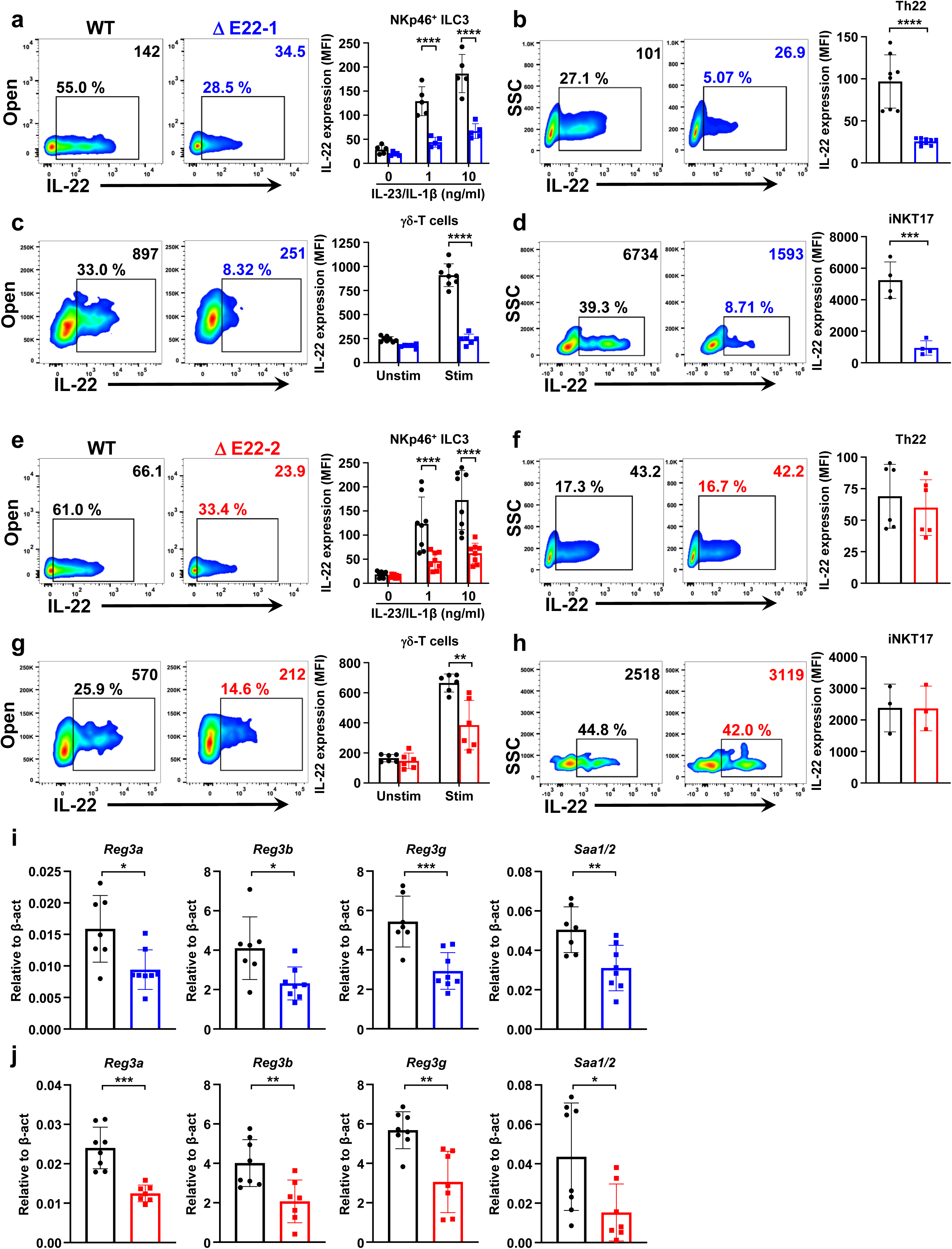
Differential regulation of IL-22 by E22-1 and E22-2 in innate vs adaptive type 3 immune cells. (a-d) Representative flow plots (left) and quantification of IL-22 (right, MFI) in NKp46^+^ ILC3s isolated from SI-LP after 3.5 h of ex vivo stimulation with indicated concentrations of IL-1β+IL-23 (a), ex vivo polarized Th22 (b), γδ-T cells from SI-LP (c), iNKT17 cells from lymph nodes (d) of WT (black) and E22-1^Δ^ mice (blue). (e-h) Representative flow plots (left) and IL-22 expression (right, MFI) in SI-LP NKp46^+^ ILC3 subset after ex vivo stimulation for 3.5 h with indicated concentrations of IL-1β+IL-23 (e), ex vivo polarized Th22 (f), SI-LP γδ-T cells (g), iNKT17 cells from lymph nodes (h) of WT (black) and E22-2^Δ^ mice (red). (i, j) RT-qPCR analysis of *Reg3a*, *Reg3b*, *Reg3g*, and *Saa1/2* in terminal ileum from WT (black), E22-1^Δ^ (blue), or E22-2^Δ^ mice (red). Expression relative to *β-actin*. Data are shown as mean ± SD and are pooled (a-c, e-g, i, j), or are representative (d, h) of at least two independent experiments. \**P* < 0.05; \*\**P* < 0.01; \*\*\**P* < 0.001; \*\*\*\**P* < 0.0001; two-way ANOVA with Sidak’s multiple comparisons test (a, c, e, g) or unpaired t-test (b, d, f, h-j).

We next examined the E22-2^Δ^ single knockout and observed a decrease in IL-22 expression for ILC3 subsets that was comparable to reductions observed in the E22-1^Δ^ and E22-1/2^Δ^ mice (**Fig. 4e and S4m-q**). Importantly, however, IL-22 production was unaffected in Th22 and NKT17 cells from the E22-2^Δ^ animals (**Fig. 4f, 4h, S4r,s, and S4u,v**). As shown in **Fig. 4g and S4t**, the E22-2 deletion led to a modest, but significant decrease (∼2-fold) in IL-22 expression by γδ-T cells. The reduced levels of IL-22 in either E22-1^Δ^ or E22-2^Δ^ mice led to a downregulation of *Reg3*α, *Reg3*β, *Reg3*γ, and *Saa1/2* in intestinal epithelium (**Fig. 4i,j**). Taken together, E22-1 and E22-2 both contribute to optimal IL-22 production in ILC3s and γδ-T cells; whereas, E22-2 is dispensable for normal induction of this cytokine in Th22 and NKT17 cells, suggesting an ILC3 and γδ-T cell-specific function for E22-2. Surprisingly, deletion of both elements had no additional impact on IL-22 expression in any of the type 3 lymphoid subsets tested, suggesting that the two enhancers are partially compensatory rather than synergistic in ILC3s and γδ-T cells (see discussion).

In addition to its role as a protective cytokine, IL-22 has a pathologic function in inflammatory diseases, such as IBD and psoriasis. These effects are attributable, in part, to its function in inducing pro-inflammatory genes, as well as anti-apoptotic and proliferative pathways. In this regard, IL-22 contributes to the development of psoriasis-like lesions following application of the topical pro-inflammatory agent, IMQ^19^. IMQ-induced psoriasis is characterized by redness and scaling of the skin, thickening of the epidermal layer from hyperplasia and keratin retention, as well as thickening of the dermis from inflammatory infiltrates and edema. The importance of IL-22 in the pathogenesis of human psoriasis is underscored by a linkage between deficits in IL-22 binding protein, elevated levels of IL-22 in skin, and enhanced incidence of lesions^18,31^.

To test whether *Il22* enhancers with distinct activity profiles differentially affect IL-22 mediated psoriasis, we treated each of the mutant mice with topical administration of IMQ to their ears. WT mice developed IMQ-induced psoriasis-like lesions on their ears over a period of 9 days, and this pathology was reduced in each of the enhancer-deleted animals (**Fig. S5a**). As a more quantifiable measurement, we found that all of the mutant mice had reduced thickening of the pinna compared to WT mice **(Fig. 5a)**. These differences were also evident when using a modified psoriasis area and severity index (PASI) score, which qualitatively quantifies redness, scaling, and thickness of psoriasis lesions **(Fig. 5b-e)**^32^. In addition, IMQ-treated WT mice showed a significant increase in the thickness of the epidermal layer histologically compared to E22-1^Δ^, E22-2^Δ^, and E22-1/2^Δ^ mice **(Fig. 5f)**. Notably, mice lacking the innate cell-specific enhancer, E22-2, had a reduced pathology that was comparable to animals lacking the E22-1 enhancer, which is also functional in Th17/22 cells. These data suggest that IL-22 expressed by ILC3 and γδ-T cells in the skin is sufficient to drive psoriasis pathogenesis in this mouse model.

**Fig. 5:**
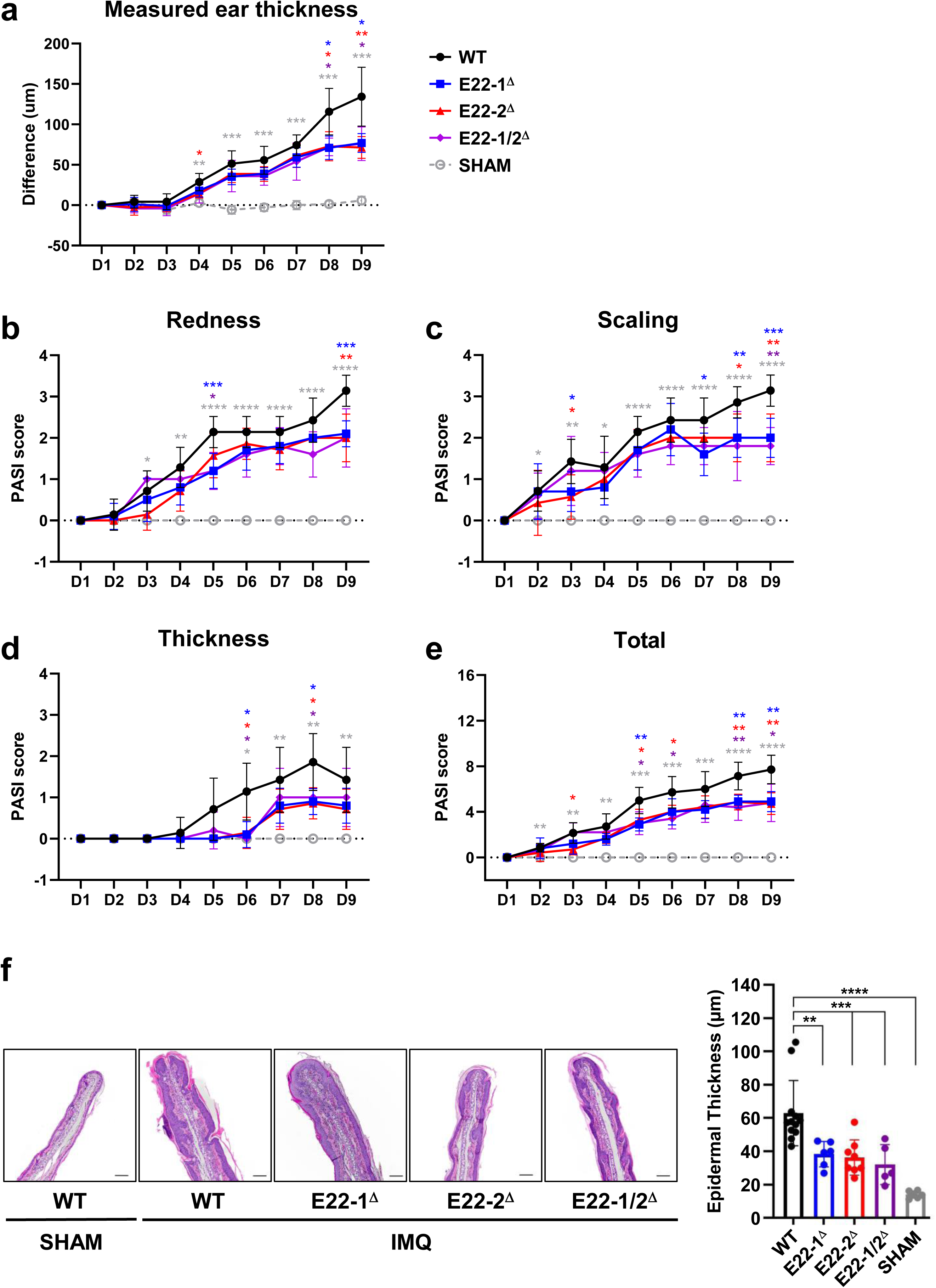
Loss of E22-1 or E22-2 ameliorates imiquimod-induced skin inflammation. Mice were treated with 31 mg of IMQ or control cream (Vaseline) on a single ear for 9 days. (a) Difference in ear measurements throughout the 9-day experiment. (b-e) Daily modified psoriasis area and severity index (PASI) scoring was used to evaluate the redness, scaling, and objective thickness of ears on a scale of 0 to 4. (f) Hematoxylin-eosin staining analysis. Representative images of the ear skin of mice (left, black bar = 100 µm) and measured thickness of epidermal layer by light microscopy (right). Data are shown as mean ± SEM and are representative of three independent experiments; n = 5-10. * P < 0.05, ** P < 0.01, *** P < 0.001, **** P < 0.0001; two-way ANOVA (a-e), or one-way ANOVA (f) followed by multiple comparisons using WT as control.

### Cell type-specific enhancer activity of E22-2

Multiple lines of evidence suggest that E22-2 is an ILC3-specific enhancer that may also have activity in a subset of γδ-T cells. To more directly test its activity profile, we analyzed a published transgenic mouse engineered to carry an IL-22-eGFP reporter that has only E22-2^33^. The BAC transgene in this mouse strain has GFP inserted upstream of the *Il22* coding sequences in exon 1, disrupting its signal peptide and ensuring cytoplasmic GFP expression (**Fig. S6a**). With regard to regulatory regions, the BAC transgene contained ∼30 kb of sequence 5’ to the *Il22* promoter. As such, the BAC reporter included E22-2, but lacked E22-1, the latter of which is required for expression of endogenous IL-22 in polarized Th cells. Indeed, the BAC reporter lacking E22-1 was silent in ex vivo polarized Th22 cells, despite expression of endogenous IL-22 in ∼20% of the cultured cells (**Fig. 6a**). In contrast, GFP was expressed in short-term cultures of primary SI-LP ILC3s (∼40% of cells), even in a fraction of cells that were not expressing endogenous IL-22 (**Fig. S6b)**. Upon stimulation with type 3 agonists, the percentage of GFP^+^ cells increased (∼60% of cells), and a higher proportion expressed endogenous IL-22 (**Fig. S6b,c**).

**Fig. 6:**
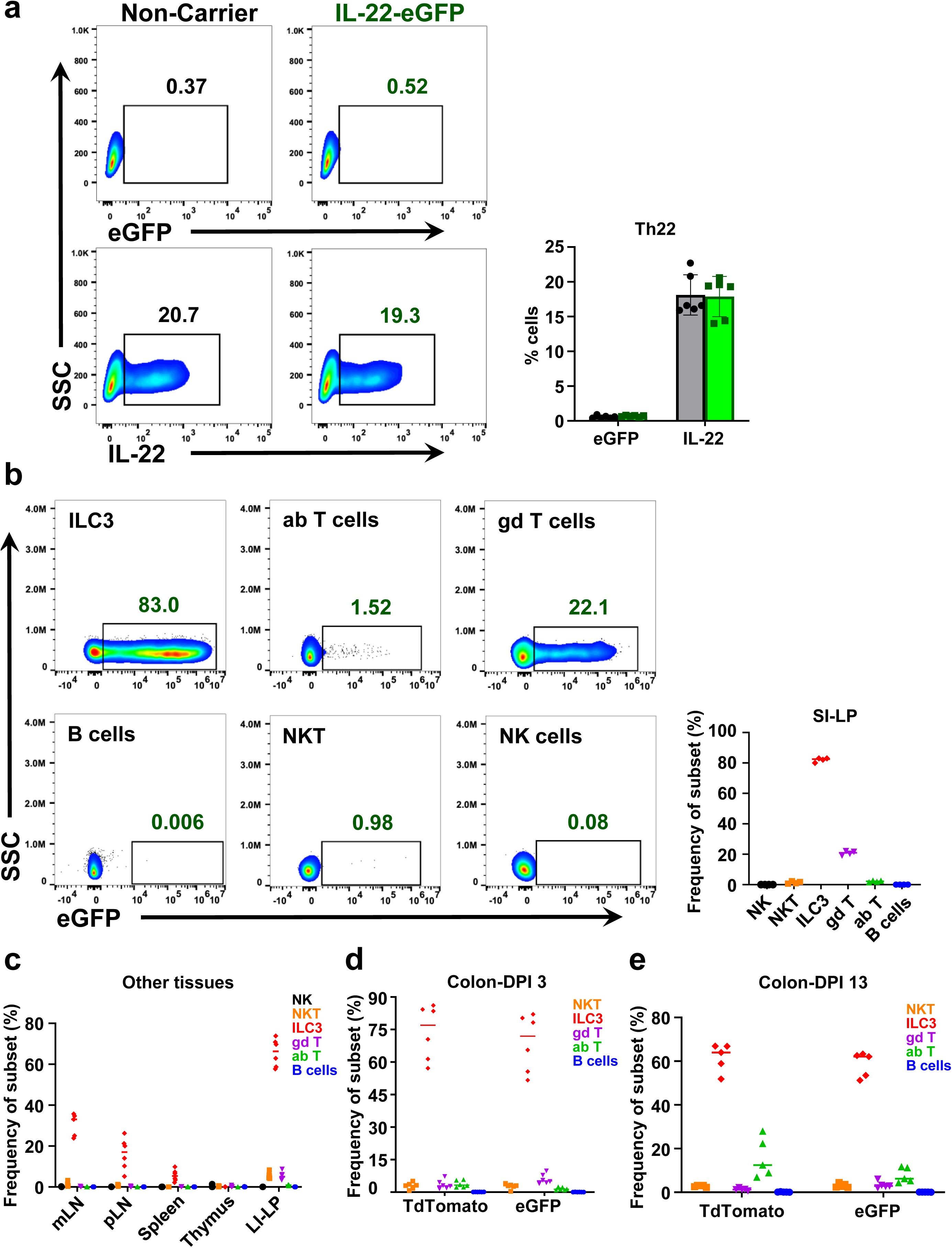
E22-2 is predominantly an ILC3 enhancer. (a) Flow cytometric analysis of reporter (eGFP) and endogenous IL-22 in ex vivo polarized Th22 cells derived from IL-22-eGFP reporter or non-carrier mice, with percentage of positive cells shown in the right (non-carrier, gray and reporter mice, green). (b) Flow cytometric analysis of reporter expression in total ILC3, αβ-T, γδ-T, B, NKT and NK cells from unfixed SI-LP isolates. Representative flow plots (left) and frequency of eGFP^+^ cells for analyzed cell types (right). (c) Flow cytometric analysis of frequency of eGFP^+^ NK, NKT, ILC3, γδ-T, αβ-T, or B cells in unfixed mLN, pLN, spleen, thymus and LI-LP isolates harvested from IL-22-eGFP reporter mice. (d,e) Flow analysis of reporters expression in NKT, ILC3, γδ-T, αβ-T, and B cells from unfixed colon-LP isolates. IL-22-eGFP X Catch22 mice were infected orally with 2x10^9^ CFU of *C. rodentium* and colon-LP were analyzed for the expression of eGFP or TdTomato, (d) 3 day post infection (DPI) or (e) 13 DPI in the indicated lymphoid subsets. Data are shown as median and pooled from at least two independent experiments.

In these initial experiments, ILC3s were fixed to stain for endogenous IL-22. However, fixation can result in loss of GFP fluorescence signal^34^. Indeed, when we directly analyzed SI-LP cell populations without fixation, 80-90% of ILC3s and ∼20% of γδ-T cells were GFP^+^, whereas the reporter was expressed in less than 2% of the other cell types examined, including αβ-T cells (**Fig. 6b**). We also examined reporter expression in cell populations from different primary or secondary lymphoid tissues. As shown in **Fig. 6c**, the E22-2-driven reporter was expressed robustly in ILC3s derived from all secondary lymphoid organs (intestinal and peripheral lymph nodes, spleen); whereas GFP was undetectable in thymocytes, NK, B, αβ-, γδ-, or NK-T cells from these tissues. In the large intestine LP (LI-LP), we observed a low percentage of GFP^+^ γδ-T and NKT cells (∼5%), but their proportion paled in comparison with the >70% of ILC3s expressing the reporter in this environment. Hence, in line with our enhancer knockout data, E22-2 is sufficient to drive gene expression in ILC3s and in a small subset of γδ-T cells, but not in conventional CD4^+^αβ-T cells.

To further establish E22-2 specificity during an immune response, we monitored LI-LP cells during *C. rodentium* infection. To also have a readout for endogenous *Il22*, we crossed the IL-22-eGFP transgene (E22-2 only) with the Catch22 strain, which harbors an IL-22 knock-in/knock-out reporter (TdTomato) and, as such, retains all *Il22* regulatory elements^35^. IL-22-eGFP x Catch22 mice were sacrificed at day 3 or day 13 post-infection and unfixed colon-LP cells were analyzed by flow cytometry. As expected, endogenous IL-22 expression, as measured by TdTomato, and expression of the E22-2-driven GFP reporter was largely limited to ILC3s at the early time point, but both were also detected in a very small percentage of γδ-T cells (**Fig. 6d**). In contrast, both reporters were silent in αβ-T cells. At the later time point, when both ILC3s and αβ-T cells are involved in the response to *C. rodentium*^13,14^, a high percentage of ILC3s expressed both reporters. Although ∼20% of αβ-T cells (**Fig. 6e**), presumably Th17/22 cells, expressed the knock-in reporter (TdTomato), <5% expressed the E22-2-driven reporter (GFP). Moreover, only a low percentage of γδ-T cells expressed either of the reporters at each time point. Collectively, these data establish that the enhancer function of E22-2 is primarily restricted to ILC3s in vivo, with modest activity in a subset of γδ-T cells but very low to no activity in a small subset of αβ-T cells. Thus, E22-2 has a rare activity profile in which its functions are restricted to the innate lymphoid counterpart of its T helper subset.

### Transcription factors required for E22-2 function and specificity

To determine the core elements of E22-1 and E22-2 that are crucial for their distinct activity profiles, we divided each of the deleted regions into segments (**Fig. S7a**) and performed luciferase reporter assays in Mnk3. E22-1 enhancer activity is restricted to a 400 bp fragment, called E22-1b, which coincides with the core of its corresponding ATAC peak in lymphoid cells (**Fig. S7b**). A prior study had shown that this region is an enhancer in cultured T cells, and it requires Runx1 and RORγt binding motifs for its activity^36^. As such, we focused on E22-2, whose activity appears to be primarily restricted to ILC3s and a small subset of gut γδ-T cells. Once again, the core ATAC peak from E22-2, which is approximately 460 bp (E22-2c), harbors all enhancer activity in Mnk3 cells (**Fig. S7b**). In silico analysis of E22-2c using the JASPAR algorithm predicted binding sites for Runx, Bcl11b, RORγt and Batf/AP-1 TFs (**Fig. 7a**). To test their individual contributions to E22-2c enhancer function, we mutated each site and performed luciferase assays. As shown in **Fig. 7b**, each of the three Runx sites, a Batf site, and a Runx-Batf overlapping site in E22-2 were absolutely required for its enhancer function in this ILC3 cell line. Given their expression patterns in Mnk3 cells (**Fig. S7c,g**) it seems likely that Batf, a TF also important for ILC3 functions,^37,38^ is responsible for the induction in enhancer activity of E22-2 following exposure to type 3 agonists.

**Fig. 7:**
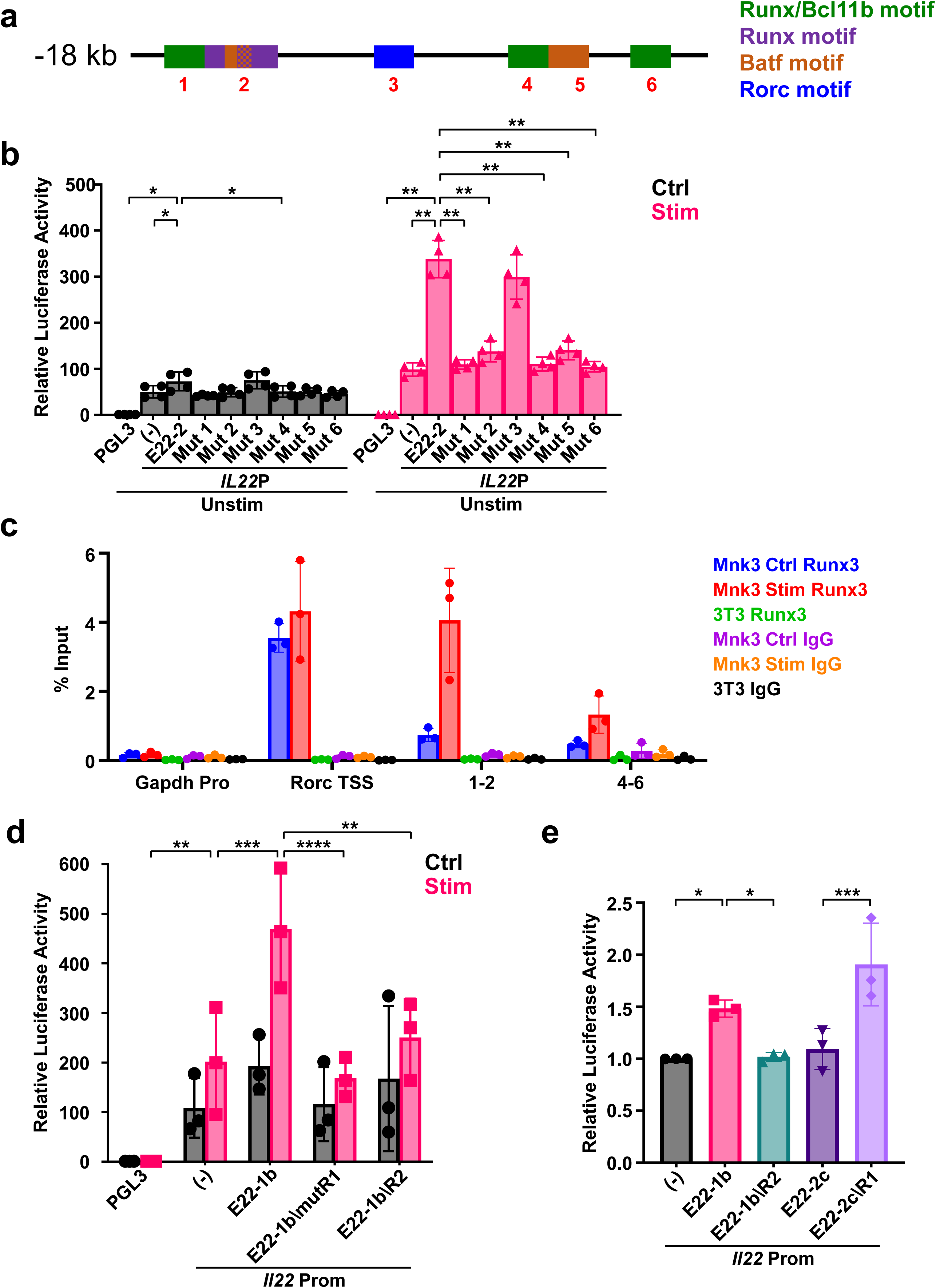
ILC3 restriction of E22-2 enhancer activity. (a) Schematic showing JASPAR predicted TF binding sites in the core E22-2 element. (b) Luciferase reporter activity of E22-2c enhancer region harboring mutations, as denoted in (a), of each binding motif. Assays were performed as in Fig. 2. (c) CUT&RUN with anti-Runx3 and anti-IgG control in Mnk3 and 3T3 cells. qPCR analysis for Runx3-bound DNA at the *Gapdh* promoter (negative control), *Rorc* promoter (positive control), sites1-2, and sites 4-6. Results are represented as percent of input chromatin. (d) Luciferase reporter activity in Mnk3 cells (+ or – IL-1β+IL-23 stimulation) of E22-1b, E22-1b with mutated RORγt motif (E22-1b/mutR1), or E22-1b harboring a replacement of its WT RORγt motif with the variant from E22-2c (E22-1b/R2). The reporter activity is relative to PGL3-Basic vector. (e) Luciferase reporter assay showing enhancer activity of E22-1b, E22-1b with the E22-2c γt motif swap (E22-1b/R2), E22-2c, or E22-2c with its WT RORγt site replaced with the E22-1b RORγt motif (E22-2c/R1) in ex vivo polarized Th22 cells. The polarized Th22 cells were derived from Catch22 reporter mice, flow sorted for TdTomato and were treated with 1X cell stimulation cocktail in the presence of anti-CD3 and anti-CD28. The reporter activity is relative to PGL3-*Il22* promoter only vector. Data are shown as mean ± SD and are pooled from four (b) or three (c-e) independent experiments. \**P* < 0.05; \*\**P* < 0.01; \*\*\**P* < 0.001; two-way ANOVA with Dunnett’s multiple comparisons test (b, d) or one-way ANOVA followed by Fisher’s test (e).

Among the members of the Runx TF family, Runx3 is necessary for ILC3 development and is expressed at the highest level in Mnk3 (**Fig. S7c**) and primary ILC3s^39^. To test whether this factor binds to E22-2, we performed CUT&RUN using a Runx3 antibody followed by qPCR analysis. We obtained robust signals for Runx3 binding in Mnk3 cells at its known site in the *Rorc* promoter but detected no binding at a negative control site in the *Gapdh* promoter (**Fig. 7c**)^39^. Importantly, Runx3 binding was detected at both ends of E22-2c, which contain sites 1-2 or sites 4-6, and these signals were more intense upon stimulation of Mnk3 cells with type 3 cytokines. In contrast, no signals were detected at any of these sites in 3T3 fibroblasts, which do not express Runx3 (**Fig. 7c**). Taken together, Runx3 binding is essential for E22-2 enhancer activity in this ILC3 cell model.

We next tested whether, similar to E22-1, the predicted RORγt site was important for E22-2 enhancer activity^36^. Analysis of published RORγt Chip-seq data from Th17 cells^40^ revealed strong binding of this factor at E22-1 but minimal binding at E22-2 (**Fig. S7d)**. Indeed, mutation of this site in E22-2 had no significant impact on reporter expression in Mnk3 cells (**Fig. 7b**). Given that the RORγt site is functional in E22-1 but not in E22-2, we compared their sequences and found that the E22-1 site was a better match to the JASPAR consensus motif (**Fig. S7e**). To determine the impact of the variant motifs on enhancer function, we first confirmed that mutation of the RORγt motif in E22-1b resulted in a loss of reporter activity in the Mnk3 line (**Fig. 7d**). In addition, we swapped the RORγt site in E22-1b with the potentially inactive site from E22-2c. As shown in **Fig. 7d**, the motif swap inactivated E22-1b, confirming that the E22-2c site cannot support enhancer function in Mnk3 cells, which express the *Rorc* gene. Importantly, we also performed the complementary motif swap and found that replacement of the RORγt site in E22-2c with its counterpart from E22-1b endowed this chimeric enhancer with activity in primary, activated Th22 cells isolated as TdTomato^+^ from Catch22 mice (**Fig. 7e**). Likewise, this replacement also augmented E22-2c enhancer function in Mnk3 cells (**Fig. S7f**), further underscoring the superior binding capability of the RORγt site from E22-1. We conclude that rather than having an ILC3-specific activation mechanism, E22-2 lacks a functional RORγt binding site, which is required for its enhancer activity in adaptive T cells. This enhancer activity profile – functional only in innate lymphoid cells – is rare among regulatory elements that control the expression of genes in Th-ILC counterparts.

## Discussion

Cytokine gene expression is regulated at multiple levels to strike a delicate balance between homeostatic fitness, productive immune responses during infection or injury, and avoiding pathological consequences of persistent overexpression, such as chronic inflammation. In this study, we examined regulatory programs for the signature type 3 cytokine, IL-22, which is expressed at the steady state in barrier tissues, primarily by ILC3s, to control homeostatic programs of immune defense and tissue integrity. Upon infection or injury, IL-22 expression is induced in innate and adaptive type 3 lymphoid cells via signals emanating from antigen or cytokine receptors. As observed during infection by the gut pathogen, *C. rodentium*, IL-22 regulatory programs permit rapid cytokine deployment by ILC3s to control damage and pathogen expansion until Th17/22 cells are fully activated to clear the pathogen^13,14,41^. Although the temporal, cell type- and agonist-specific control of IL-22 has been appreciated for some time, the cis-elements responsible for these important regulatory features remained largely unknown^42^. Indeed, most transcriptional control elements for genes involved in polarized immune responses, including type 3 responses, are equally functional in Th and ILC counterparts.

Here, we performed a comprehensive CRISPRi screen for cis-elements that regulate IL-22 expression in type 3 lymphoid cells. We discovered two enhancers that, together, mediated IL-22 expression for homeostatic defense in the gut (i.e., AMP expression), as well as protection from *C. rodentium* infection. Importantly, both enhancers also contributed to pathologic IL-22 expression during skin inflammation in a mouse model of psoriasis. The activity profile of one enhancer, E22-1, resembled that of others involved in Th-ILC responses, in that it was required for normal levels of IL-22 in both innate and adaptive immune cell counterparts. In contrast, E22-2 exhibited a rare, nearly exclusive activity profile in ILC3s, driving IL-22 expression in these cells at steady state, following induction with type 3 agonists, and during *C. rodentium* infection. Although E22-2 could support reporter expression in a subset of γδ-T cells, overall levels were at least a magnitude lower compared with ILC3s. Thus, E22-2, which is highly conserved between humans and mice, has evolved to be an innate lymphoid-specific enhancer for IL-22 expression in barrier tissues.

Given its unique activity profile among other known Th-ILC enhancers, the purpose for an ILC3-specific control element for the *Il22* gene remains uncertain. One possible explanation is the need for steady state expression of IL-22 by ILC3s to support homeostatic levels of AMPs and barrier fitness. This may require the enhancer to have a weaker constitutive activity but remain inducible to rapidly achieve higher levels of IL-22 when tissue-resident ILC3s receive cues resulting from perturbations of homeostasis. Indeed, compared with E22-1, the ILC3-specific E22-2 drives steady state expression of a reporter gene, but appears to be a weak constitutive enhancer in cell models. However, the enhancer is highly inducible in all three ILC3 subsets, when the cells are exposed to the type 3 agonists, IL-23 and IL-1β. By contrast, the Th22/ILC3 enhancer, E22-1, does not appear to be required for basal expression of the IL-22 reporter in vivo but clearly contributes to inducible expression of the cytokine in all type 3 cells ex vivo. Together, the distinct activity profiles of the two enhancers likely provide flexibility for IL-22 expression that is suited for balancing steady state fitness, rapid antigen-independent innate responses, and delayed antigen-specific adaptive responses to pathogens.

Although E22-1 appears to be the dominant enhancer for inducible IL-22 expression in CD4^+^ T cells, both enhancers are required for inducible expression of the cytokine in ILC3s and a subset of γδ-T cells. Surprisingly, however, the activities of E22-1 and E22-2 in ILC3s were not additive; mice harboring deletions of both enhancers phenocopied those of either single deletion in stimulated ILC3s, albeit all had residual levels of IL-22 expression over baseline. Although details remain to be resolved, these data suggest that the enhancers are either partially compensatory in activated ILC3s or perhaps, a contingent mechanism is at play, whereby E22-2 is required for full activation of E22-1 in ILC3s. However, one cannot rule out that the two enhancers have an additive function under certain conditions in vivo, especially given the technical difficulties associated with quantifying IL-22 expression in situ. We also cannot rule out that additional *Il22* regulatory elements escaped our CRISPRi screen, such as in the super-enhancer region, due to functional redundancies in the Mnk3 cell model, a possibility that could explain low level residual IL-22 expression in the enhancer double knockouts.

The exceptional activity profile of E22-2 distinguishes it from E22-1 and other regulatory elements associated with genes expressed in Th-ILC counterparts. Mechanistically, both of the *Il22* enhancers depend on binding sites for Runx TFs^36^ (this study). For E22-2, these sites appear to be bound by the ILC3 dominant family member, Runx3, in an ILC3 cell model. Unlike E22-1, the ILC3-specific E22-2 has multiple Runx sites, each of which are absolutely required for its enhancer activity, perhaps endowing it with its modest basal function. Importantly, E22-2 lacks a functional RORγt binding site, which appears to be the only restriction for it not having enhancer activity in T cells, as shown by replacing its non-consensus site with the functional RORγt motif from E22-1. Since RORγt is likely activated in all type 3 cells, this may explain the more promiscuous activity of E22-1, while having only Runx sites restricts E22-2 to innate cells. Strikingly, JASPAR analysis of human E22-1 and E22-2 revealed retention of these transcription factor motif layouts. Future studies may identify this architecture – multiple Runx sites, no functional RORγt sites – in additional ILC3-specific enhancers, although the reason for such cell-type specificity remains to be further clarified. Notwithstanding, the restricted activity of E22-2 to ILC3s may provide a means to generate a much-needed tool: an ILC3-specific Cre driver in mice, which would be invaluable for dissecting gene functions in Th22 versus ILC3 counterparts.

## Methods

### Mice

All mice used in the study were on the C57BL/6 background. E22-1^Δ^, E22-2^Δ^, and E22-1/2^Δ^ mice were generated using CRISPR-Cas9 technology at the transgenic core of Children’s Hospital of Philadelphia Research Institute and the deletions in founder mice were confirmed by sequencing. The sgRNAs (Table S1) used were designed by the GEiC core at Washington University in St. Louis. Catch22 mice^35^ were kindly provided by Richard M. Locksley and IL22-eGFP transgenic mice^33^ were purchased from the Jackson Laboratory (035005, Jackson Laboratory). For some experiments, Catch22 and IL22-eGFP mice were intercrossed to generate Catch22 x IL22-eGFP mice. All animals were bred and maintained in specific pathogen-free animal facility at The Ohio State University. Mice used for experiments were sex- and age-matched. All experiments and procedures were performed in accordance with IACUC guidelines.

### Cell lines and culture

EL4 and 63-12 cells were cultured in RPMI (Gibco) supplemented with 10% fetal bovine serum (Sigma), 1X Penicillin-Streptomycin (Gibco), and 50 µM β-mercaptoethanol (Sigma). Mnk3 cells, described previously^25^, were cultured in DMEM supplemented with 10% fetal bovine serum, 1X Penicillin-Streptomycin, 50 µM β-mercaptoethanol, 10 ng/ml mouse IL-7 (R&D Systems, 407- ML/CF), and 10 ng/ml mouse IL-2 (R&D Systems, 402-ML/CF). HEK293T cells were maintained in DMEM (Gibco) supplemented with 10% fetal bovine serum, 1X Penicillin-Streptomycin, and 50 µM β-mercaptoethanol. For lentivirus production, HEK293T cells were co-transfected with lentiviral (1 µg), psPAX2 (1 µg), and pMD2.G (0.125 µg) plasmids using the TransIT-293 transfection reagent (Mirus, MIR 2700). Virus particles were harvested ∼48 hours post-transfection and passed through a 0.45 µm syringe filter. Mnk3 cells were transduced with lentiviral supernatant in the presence of 5 µg/ml polybrene (Millipore Sigma, TR-1003-G) For experiments where sgOpti lentiviral particles were used, cells were selected for 72 hours with puromycin (1 µg/ml) after ∼48 hours of transduction.

For the generation of Mnk3i lines, Mnk3 cells were first transduced with rtTA3 lentiviral particles (pLenti CMV rtTA3 Blast, Addgene Plasmid #26429) and selected using blasticidin (10 µg/ml) after 48 hours of viral infection. After checking the expression of rtTA3 in blasticidin selected cells, we next infected them with TRE3G-dCas9-KRAB-P2A-mCherry lentiviral particles. After 48 hours of infection, cells were treated with dox (2 µg/ml) for 24 hours, flow sorted for mcherry high cells and cloned by limiting dilution. The single cell clonal populations were screened, and a clone was selected with highest mCherry expression, a surrogate for high dCas9-KRAB expression.

### Cloning

For Lenti-TRE3G-dCas9-KRAB-P2A-mCherry construct, tetracycline inducible TRE3G promoter was amplified from c3GIC9 (Addgene Plasmid #62191) vector adding EcoR1 restriction site at 5’ end and Mlu1 followed by BamH1 restriction enzyme sites at the 3’ end. Next, TRE3G promoter and pHR-pEF-EGFP vector (Addgene Plasmid #67952) were digested with EcoR1+BamH1 and ligated, resulting in pHR-TRE3G-EGFP vector. Lastly, dCas9-KRAB-P2A-mcherry cassette was swapped from pHR-SFFV-KRAB-dCas9-P2A-mCherry (Addgene Plasmid #60954) into pHR-TRE3G-EGFP vector from above using Mlu1 and Not1 restriction sites. sgRNAs targeting *Il22* promoter, E22-1, E22-2, twin-E22-2, or scrambled sgRNA were cloned downstream of the U6 promoter in sgOpti (Addgene Plasmid #85681).

### ATAC-seq

Nuclei were isolated from freshly harvested cells (∼6x10^4^) using ATAC lysis buffer (10 mM Tris-HCL pH 7.4, 10 mM NaCl, 3mM MgCl2, and 0.1% IGEPAL CA-630). The nuclei were then subjected to transposition reactions for 30 minutes using Tagment DNA TDE1 Enzyme (Illumina, 20034197). Purified DNA was then amplified to generate libraries using Nextera XT DNA Library Preparation Kit (Illumina, FC-131-2001). The libraries were quantified, assessed for quality and paired-end sequencing was performed with Illumina HiSeq.

### CRISPRi screen

The CRISPRi screen strategy was adapted from a previously published study^43^. ATAC- and H3K27ac-seq data from Mnk3, Mnk1, NK, ILC1, Th1, ILC3, and Th17 cells (new and published data^26,27^) were analyzed to identify all active/accessible regions in a 1.5 Mb space around the *Il22* mouse locus (chr10:117149900-118685961, mm9 genome build). Each of the selected regions were then extended to include ±200 bp (Table S2). The library pool was designed and cloned into sgOpti plasmid by GEiC core at Washington University. Mnk3 cells were derived from NIH-Swiss mouse, so we performed whole genome sequencing for this cell line to identify the variants when compared with the sequence of C57BL/6 genome (used for designing sgRNAs). The pool appeared to be equally effective in either mouse background. The pool consisted of 20-25 sgRNAs per regulatory region, positive controls (e.g., *Il22* promoter), negative controls (ATAC negative region in chromosome 10 and promoters of certain non-essential genes such as CXCR4), and published non-targeting controls for mouse^44^ (Tabel S2). ∼82% of the above targets were covered by ≥20 sgRNAs while the remaining had fewer sgRNAs. The Mnk3i line was infected with the lentiviral library at an MOI<0.3. After 48 hours of transduction, cells were subjected to 72 hours of puromycin (1 µg/ml) selection. The selected cells were then either treated with dox (2 µg/ml) or left untreated for 24 hours and then incubated with 10 ng/ml each of IL-1β and IL-23 for additional 16 hours. Cytokine-stimulated cells were treated with GolgiStop for 6 hours and stained intracellularly with IL-22 and IL-17F antibodies. ∼21x10^6^ IL-17F^+^ cells (700-fold coverage of library) were flow sorted into IL-22-negative, -low, -medium, and -high fractions. Genomic DNA was extracted from the cellular fractions, and integrated sgRNA sequences were amplified from 19 µg genomic DNA (250-fold coverage of library) of each sample using NEBNext Ultra II Q5 (NEB, M0544S) using custom designed oligos (Table S2). The libraries were sequenced using custom sequencing primers with Illumina HiSeq 2500. The data were analyzed using MAGeCK v0.5.7.^28^ Normalized count for each sample was calculated (Table S3) and sgRNA enrichment or depletion in +dox sorted cellular populations was assessed by comparing their sgRNA abundance with corresponding -dox sorted cellular fraction.

### Luciferase assay

The *Il22* promoter was inserted into NheI and XhoI sites present upstream from luciferase gene of PGL3-Basic vector. The enhancer elements E22-1, E22-2, E22-1a, E22-1b, E22-1c, E22-2a, E22-2b, E22-2c, or E22-2d were cloned downstream of luciferase gene and the SV40 poly(A) signal in the PGL3-*Il22* promoter construct. Mutants were created using Q5 Site-Directed Mutagenesis Kit (NEB, E0554) after designing primers (Table S1) with the NEBaseChanger tool. Cells were resuspended in Chica buffer (5 mM KCl, 15 mM MgCl2, 0.12 M Phosphate Buffer pH 7.2, 25 MM Sodium succinate, and 25 mM Manitol) and co-nucleofected with the 5 µg of firefly luciferase and 100 ng of renilla constructs (50:1 ratio). Mnk3 cells were nucelofected using program X-005 of Nucleofector^®^ 2b System, rested for 24 hours and then either left untreated or stimulated with 10 ng/ml each of IL-1β and IL-23 overnight. EL4 cells were nucleofected using the EL4 program C-009, rested overnight and stimulated with Cell Stimulation Cocktail (eBioscience, 00-4970-93) for 4 hours. For 63-12 cells, the X-001 program was used for nucleofection. Harvested cells were assayed in a 96-well plate using Dual-Glo® Luciferase Assay System (Promega; E2920) following the manufacturer’s instructions.

Splenic naïve CD4^+^ T cells from Catch22 mice were ex vivo polarized into Th22 cells and, on day 3, TdTomato positive cells were flow sorted and rested overnight in the presence of IL-2 (20 ng/ml), IL-6 (10 ng/ml) and IL-23 (10 ng/ml). The cells were harvested, washed with 1X PBS, resuspended in P4 buffer (Lonza, V4XP-4024) and nucleofected with firefly+renilla luciferase vectors using program CM137 of 4D-Nucleofector system. The nucleofected cells were then plated in anti-CD3 and -CD28 coated plates with complete RPMI media containing IL-2 (20 ng/ml), IL-6 (10 ng/ml) and IL-23 (10 ng/ml) overnight and assayed for firefly and renilla signals as described above.

### Quantitative RT-PCR (qPCR)

RNA was extracted from cells using TRIzol and purified. Terminal ileum (∼4 cm) was cut open, rinsed with HEPES/HBSS buffer (1X HBSS, and 15 mM HEPES) and homogenized in TRIzol using soft tissue homogenizing CK14 precellys lysing kit (Bertin Technologies, P000912-LYSK1-A.0). The homogenized sample was transferred to a 1.5 ml tube and chloroform added. The sample was mixed by vigorous inversion for 15 secs, incubated at RT for 3 mins and spun @ 12000 g for 15 mins at 4°C. The aqueous layer was transferred to columns provided with Qiagen RNA isolation Kit and proceeded for RNA isolation following manufacturer’s protocol. The isolated RNA was then reverse transcribed to cDNA using M-MuLV Reverse Transcriptase (NEB, M0253) according to the manufacturer’s protocol. qPCR was performed using SYBR Green mastermix, and gene expression was normalized to that of *β-actin*. Primers used are listed in Table S1.

### Flow cytometry

The Mnk3 lines were treated with 10 ng/ml each of IL-1β and IL-23 or left untreated overnight followed by addition of GolgiStop (BD Biosciences, 554724) for 6 hours. Cells were then intracellularly stained for IL-22 (clone 1H8PWSR; eBioscience, 12-7221-82) and IL-17F (clone eBio18F10; eBioscience, 53-7471-82) using the BD Cytofix/Cytoperm™ Fixation/Permeabilization Kit (BD Biosciences, 554714) following manufacturer’s protocol. Samples were acquired on Cytek Aurora or BD flow cytometers and analyzed using FlowJo software.

### Tissue harvest and population analysis

#### Gut Lamina Propria

Small-intestines (SI) or colons were harvested and flushed to remove luminal content. Intestines were then cut open, minced into pieces and rotated in HBSS containing EDTA, FBS, and HEPES for 20 minutes, vortexed briefly and then rotated again in fresh buffer for an additional 20 minutes to remove epithelial layers, and washed thoroughly to rinse off EDTA. The tissues were then digested in complete RPMI containing Collagenase IV (100 U/ml) for 40 mins at 37°C. The digested tissues were passed through a 100 µm cell strainer and pelleted. LP lymphocytes were isolated using Percoll (Cytiva, 17089101) density gradient and washed with media. For analysis of IL-22 production by SI-LP lymphocytes, the cells were treated with either 1 ng/ml or 10 ng/ml each of IL-1β and IL-23 or left untreated followed by addition of GolgiStop after 30 minutes and further incubated for 3 hours. The cells were then stained for flow cytometry analysis. SI-LP ILC3s were identified as live CD3^-^CD5^-^CD19^-^CD45^int^CD90.2^hi^ lymphocytes and in some experiments were then further subdivided into NKp46^+^ CCR6^-^, NKp46^-^CCR6^+^, or double negative subsets. SI-LP γδ-T cells were identified as live CD19^-^CD45^+^CD3^+^TCRβ^-^TCRγ/δ^+^ lymphocytes.

#### NKT17 cells analysis

Tissue culture plates were coated for 2 hours at 37°C with anti-hamster IgG (1:50 dilution) (MP Biomedicals, 856984) followed by coating with anti-CD3 (0.25 µg/ml) (BioLegend, 100340) and anti-CD28 (1 µg/ml) (BioLegend, 102116) for 1 hour at 37°C. Cells from all peripheral (mammary, brachial, axillary, and inguinal) lymph nodes were pooled, RBCs were lysed using ACK buffer, and were cultured overnight in the above coated plates in the presence of IL-1β (10 ng/ml) and IL-23 (10 ng/ml) followed by addition of GolgiStop for additional 4 hours. Cells were first incubated with mouse CD1d PBS-57 BV421-labeled tetramer (1:100, kindly provided by NIH Tetramer Core Facility at Emory University) for 30 minutes on ice, washed twice, followed by staining for surface receptors, RORγt and cytokines (IL-22 and IL-17A) using the Foxp3 Transcription Factor Fixation/Permeabilization kit (eBioscience, 00-5521-00). NKT17 cells were identified as live lymphocytes sized CD45^+^CD3^+^TCRβ^+^RORγt^+^CD1d-PBS-57 tetramer^+^ population.

#### Ex vivo T cell polarization

Tissue culture plates were coated with anti-hamster IgG and anti-CD3+anti-CD28 as described above. Splenic naïve CD4^+^ T cells were MACS sorted using the naive CD4^+^ T Cell Isolation Kit (Miltenyi Biotec, 130-104-453) and cultured in anti-CD3/-CD28 coated plates in RPMI containing 10% FBS, 1X Glutamax, 1X Pyruvate, 1X Penicillin-Streptomycin and 50 µM β-mercaptoethanol for 3 days in Th22 polarizing conditions: IL-6 (20 ng/ml), IL-23 (20 ng/ml), FICZ (200 nM), 100 ng/ml each of anti-IL-2, anti-IL-4, anti-TGFβ1,2,3 and anti-IFNγ. For cytokine analysis, cells were stimulated with 1X Cell Stimulation Cocktail (plus protein transport inhibitors) (eBioscience, 00-4975-03) for 4 hours and stained intracellularly using IL-22 (Clone Poly5164; BioLegend, 516404) and IL-17A (Clone TC11-18H10; BD Biosciences, 560220) antibodies for flow cytometry analysis.

LIVE/DEAD™ Fixable Aqua Dead Cell Stain (Molecular Probes Invitrogen, L34966) or LIVE/DEAD™ Fixable Blue Dead Cell Stain (Molecular Probes Invitrogen, L34962) was used at 1:500 dilution to identify and exclude dead cells from the analyses.

In reporter mice analysis, cell types were identified among live lymphocytes as ILC3s: CD3^-^CD19^-^ CD45^int^CD90.2^hi^CD127^+^, B cells: CD45^+^CD19^+^, NKT17 cells: CD19^-^CD45^+^CD3^+^CD1d-PBS-57 tetramer^+^, αβ-T cells: CD19^-^CD45^+^CD3^+^TCRβ^+^TCRγδ^-^CD4^+^, γδ-T cells: CD19^-^ CD45^+^CD3^+^TCRβ^-^TCRγδ^+^, and NK cells: CD3^-^CD19^-^CD45^+^CD127^-^NK1.1^+^.

### Antibodies

The primary mouse cells were stained using fluorochrome conjugated antibodies against CD5 (clone 53-7.3; BioLegend, 100624), CD3 (Clone 17A2 and 145-2C11; BioLegend), CD19 (Clone 1D3/CD19; BioLegend), CD45 (Clone 30-F11; BioLegend), CD90.2 (Clone 53-2.1; BD Biosciences), CCR6 (Clone 140706; BD Biosciences, 564736), NKp46 (Clone 29A1.4; BioLegend, 137616), Streptavidin PE-Cyanine7 (eBioscience, 25-4317-82), IL-22 (Clone Poly5164; BioLegend, 516404), IL-17A (Clone TC11-18H10; BD Biosciences, 560220), CD4 (Clone RM4-5; BioLegend), TCRβ (Clone H57-597; BioLegend), TCRγδ (Clone GL3; BioLegend), CD127 (Clone A7R34; BioLegend, 135043), RORγt (Clone AFKJS-9, eBiosciences, 17-6988-82), and NK1.1 (Clone PK136; BioLegend, 108707).

### Citrobacter rodentium infection

After 1 week of acclimatization, 8-12 weeks age- and sex-matched mice were administered orally with 2x10^9^ CFU of kanamycin resistant *C. rodentium* (DBS120 Kanr). The mice were observed and weighed daily for 20 days after infection. Fecal samples were collected, weighed, homogenized in 1X PBS and serially diluted. Different dilutions were plated on LB plates containing kanamycin. Colonies were counted after ∼20-24 hours of incubation at 37°C and CFU determined as CFU/g stool = (colony count X Total volume of the sample) / (Volume plated X Dilution Factor X Wt. of the stool sample in grams).

### Imiquimod-induced skin inflammation

Mice of each genotype were cohoused at least 7 days prior to the start of the experiment. The left pinnae of mice were treated topically with 31 mg of 5% imiquimod cream or a control cream (Vaseline) daily for 9 consecutive days. Fisherbrand Traceable Digital Calipers were used to measure ear thickness daily. The PASI clinical scoring system was modified from previously published report^32^. Briefly, a visual exam of the ears was performed daily to assess the redness, scaling, and thickness on a scale of 0-4 (0-none, 1-mild, 2-moderate, 3-marked, 4-severe) and the mean for each genotype was calculated. Mice were sacrificed on day 9 and the treated ear was fixed in 4% v/v paraformaldehyde for a minimum of 24 hours followed by paraffin embedding. Five µm sections were cut with a microtome and stained with hematoxylin and eosin by the Comparative Pathology & Digital Imaging Shared Resource at The Ohio State University. Samples were then observed under light microscopy and three measurements of the epidermal thickness were taken using cellSens by Olympus LS.

### CUT&RUN

CUT&RUN was performed as described^45^. Briefly, 1x10^6^ unfixed cells bound to activated ConA-coated magnetic beads (Bangs Laboratories BP531) were incubated overnight at 4°C with either 1:100 dilution of Runx3 monoclonal Antibody (Clone 2B3; Invitrogen, MA5-17169) or mouse IgG control antibody (Diagenode, C15400001-100) in antibody buffer (20 mM HEPES pH7.5, 150 mM NaCl, 0.5 mM Spermidine, protease inhibitor, 0.025% digitonin, 2 mM EDTA) followed by 2X wash with digitonin buffer (20 mM HEPES pH7.5, 150 mM NaCl, 0.5 mM Spermidine, protease inhibitor, 0.025% digitonin) and binding of protein A/G-MNase for 1 hour at 4°C. After 2X washes with digitonin buffer, beads were washed in ice-cold low salt wash buffer (20 mM HEPES, pH 7.5, 0.5 mM spermidine, 0.025% digitonin) and digested using MNase digestion buffer (3.5 mM HEPES pH 7.5, 10 mM CaCl2, 0.025% digitonin) for 5 minutes on ice. The CUT&RUN fragments were released by using stop buffer (170 mM NaCl, 20 mM EGTA, 0.025% Digitonin, 25 µg/ml glycogen, 50 µg/ml RNase A) and incubating for 30 minutes at 37°C. Input samples were prepared by fragmenting the chromatin of 1x10^6^ cells using bioruptor. The chromatin was precipitated, purified using phenol chloroform extraction method and Qiagen MaXtract tubes. Extracted chromatin was analyzed by qPCR and enrichment in each sample was reported as the signal relative to the total amount of input DNA. Primers used are listed in Table S1.

### Statistical methods

Prism (GraphPad Software) was used to perform statistical analysis and to plot the data. The statistical tests used, and p values are indicated in the figure legends and figures. P values were calculated using either unpaired t-tests, one-way ANOVA, two-way ANOVA, or three-way ANOVA.

### Data Availability

ATAC-seq, H3K27ac Chip-seq, and RNA-seq data have been deposited to NCBI Sequence Read Archive (SRA) repository under accession PRJNA1217908.

## Acknowledgments

We thank the NIH Tetramer Core Facility (contract number 75N93020D00005) for the mouse CD1d PBS-57 BV421-labeled tetramer, the Genome Technology Access Center at Washington University School of Medicine for help with genomic analyses, and the Comparative Pathology & Digital Imaging Shared Resource at The Ohio State University for processing of fixed tissues for histology. This work is supported by NIH grants AI134035 (EMO, MC), AI182416 (EMO), T32 AI165391 (LSH), DK126969, DK132327, DK30292 (MC), and a Pelotonia Foundation Postdoctoral Fellowship (AS).

## Author contributions

E.M.O., M.C., and A.S. conceptualized and supervised the study. A.S. performed most experiments, with additional contributions by L.S.H., V.A.S., M.V.D.M., N.G.S., B.B., and J.L.F. Mutant mice were generated with help from R.G. and C.H.B. Data analyses were performed by A.S., L.S.H., M.V.D.M., N.G.S., B.B., E.M.O., M.C., and K.E.H. The manuscript was written by A.S. and E.M.O. with editing by all authors.

## Supplemental method

The modified psoriasis area and severity index scoring criteria

**Table S1 (Separate file)**
|  | Redness | Scaling | Thickness |
| --- | --- | --- | --- |
| 0 | No observable redness | No observable scaling | No observable thickening |
| 1 | Mild redness at base of pinna | Mild individualized scaling | Mild rounding of margins |
| 2 | Mild redness at base of pinna with congestion of vessels | Mild multifocal scaling or moderate focal patch of scaling | Moderate rounding of margins, ear still pliable |
| 3 | Moderate diffuse redness with congestion of vessels | Focal large keratin flakes | Marked thickening with reduced pliability |
| 4 | Marked diffuse red pallor | Multifocal large keratin flakes | Severe thickening with noticeable pitting, loss of pliability |

**Table S1 (Separate file)**

**Sequences of sgRNAs, cloning primers, primers for in vitro cutting assay, RT-qPCR primers, and CUT&RUN qPCR primers.**

**Table S2 (Separate file)**

**CRISPRi screen sgRNA library design and sequences**

a. Coordinates (mm9) of controls for designing sgRNAs
b. Coordinates (mm9) of regulatory regions for *Il22* locus for designing sgRNAs
c. sgRNA library pool
d. Oligos for NGS library amplification and sequencing

**Table S3 (Separate file) CRISPRi screen results**

a. Normalized counts
b. Count summary
c. Top 10 regions with sgRNAs enriched in IL-22^neg^ and depleted in IL-22^lo^, IL-22^med^ populations

---

**Supplemental Fig. 1:**
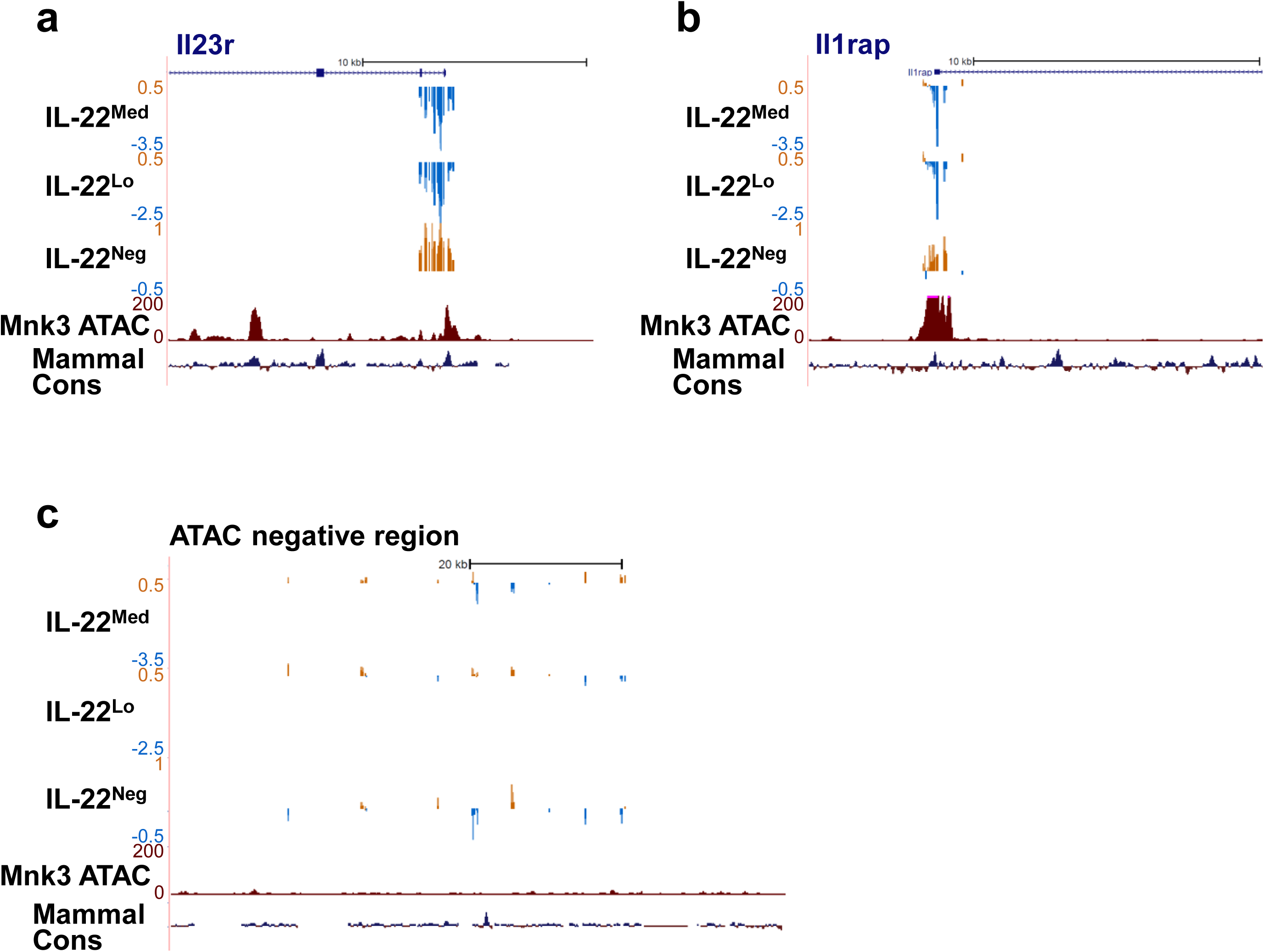
CRISPRi screen in Mnk3 cells. UCSC genome browser view showing sgRNA depletion (blue) or enrichment (gold) in the indicated sorted fractions for (a) *Il23r* promoter (positive control), (b) *Il1rap* promoter (positive control), and (c) ATAC negative region on chromosome 10 (negative control).

**Supplemental Fig. 2:**
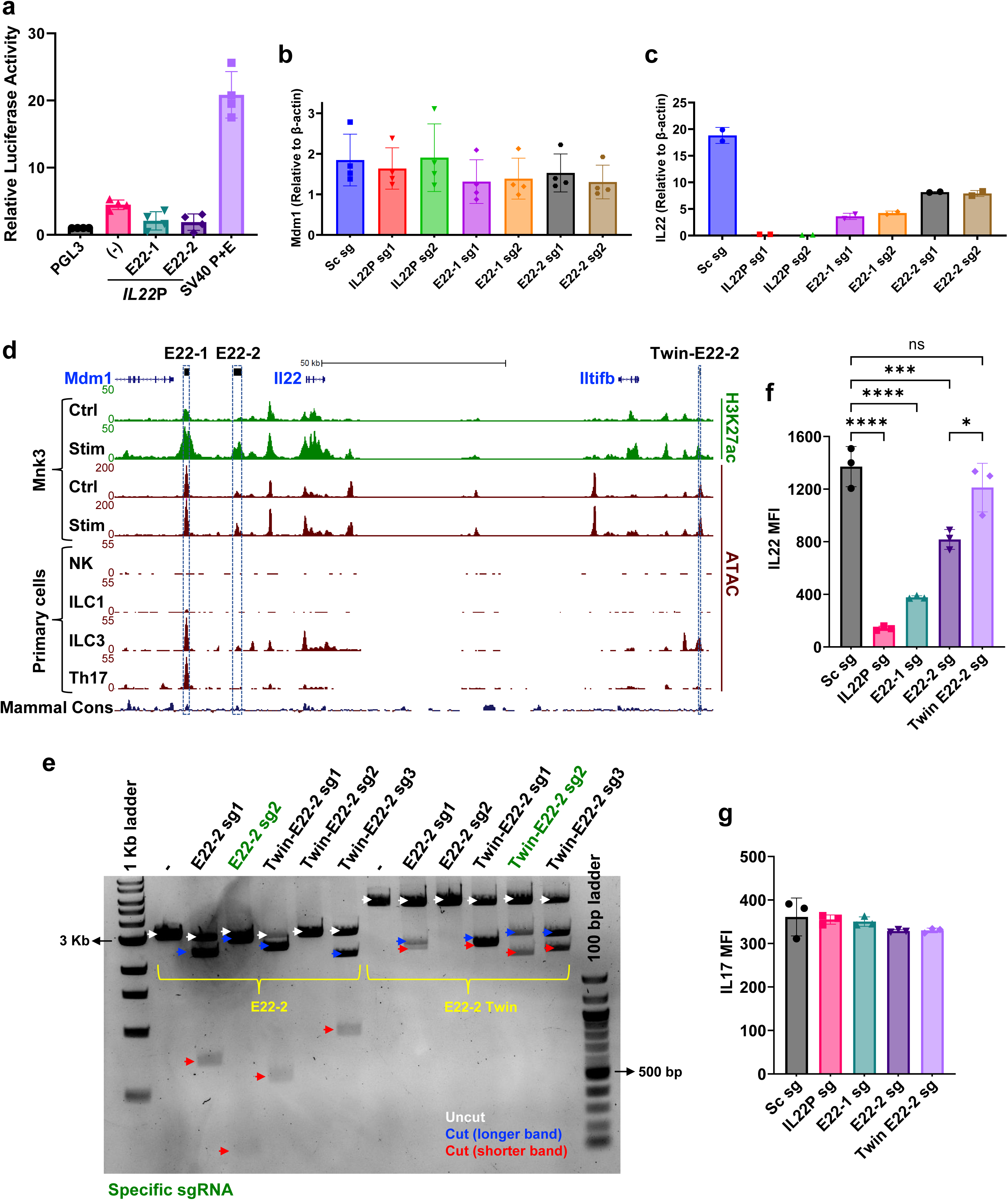
E22-1 and E22-2 act as *Il22* enhancers in Mnk3 cells. (a) Luciferase reporter assay in 63-12 cells relative to control PGL3-Basic vector. The control SV40 promoter/enhancer vector is included. (b, c) RT-qPCR analysis of (b) *mdm1* and (c) *Il22* expression relative to *β-actin*. Mnk3i lines transduced with viral particles having sgRNA targeting *Il22* promoter, E22-1, E22-2, or scrambled sgRNA were selected with puromycin, followed by 24 h of dox treatment and additional 16 h of stimulation with 10 ng/ml each of IL-1β and IL-23. (d) UCSC genome browser view of mouse *mdm1*, *Il22* and *Iltifb* genes, displaying ATAC-(maroon) and H3K27ac-seq (green) tracks. E22-1, E22-2, and twin E22-2 are highlighted in blue boxes. (e) Agarose gel analysis for in vitro cas9 cutting assay of E22-2 and twin-E22-2 with sgRNAs. E22-2 and twin-E22-2 were amplified from Mnk3 genomic DNA and purified, followed by incubation with ribonucleoprotein complexes consisting of sgRNAs targeting either E22-2 or twin-E22-2, for 1 h at 37°C. The agarose gel depicts bands of uncut (white arrows) or digested DNA (longer, blue arrows or shorter, red arrows). The sgRNAs specific for E22-2 or twin-E22-2 are labelled in green. (f, g) Flow cytometry analysis showing MFI quantification of (f) IL-22 and (g) IL-17F. Mnk3i lines transduced with sgRNAs targeting *Il22* promoter, E22-1, E22-2, twin-E22-2 or scrambled sgRNA were puromycin selected, dox treated for 24 h, followed by 16 h of stimulation with 10 ng/ml each of IL-1β and IL-23. Data are shown as mean ± SD and pooled from (a, b) four independent experiments, (c) two independent experiments, or (f, g) three independent experiments. \**P* < 0.05; \*\*\**P* < 0.001; \*\*\*\**P* < 0.0001; one-way ANOVA followed by (a, b) Dunnett’s multiple comparisons test or (f, g) Tukey’s multiple comparisons test.

**Supplemental Fig. 3:**
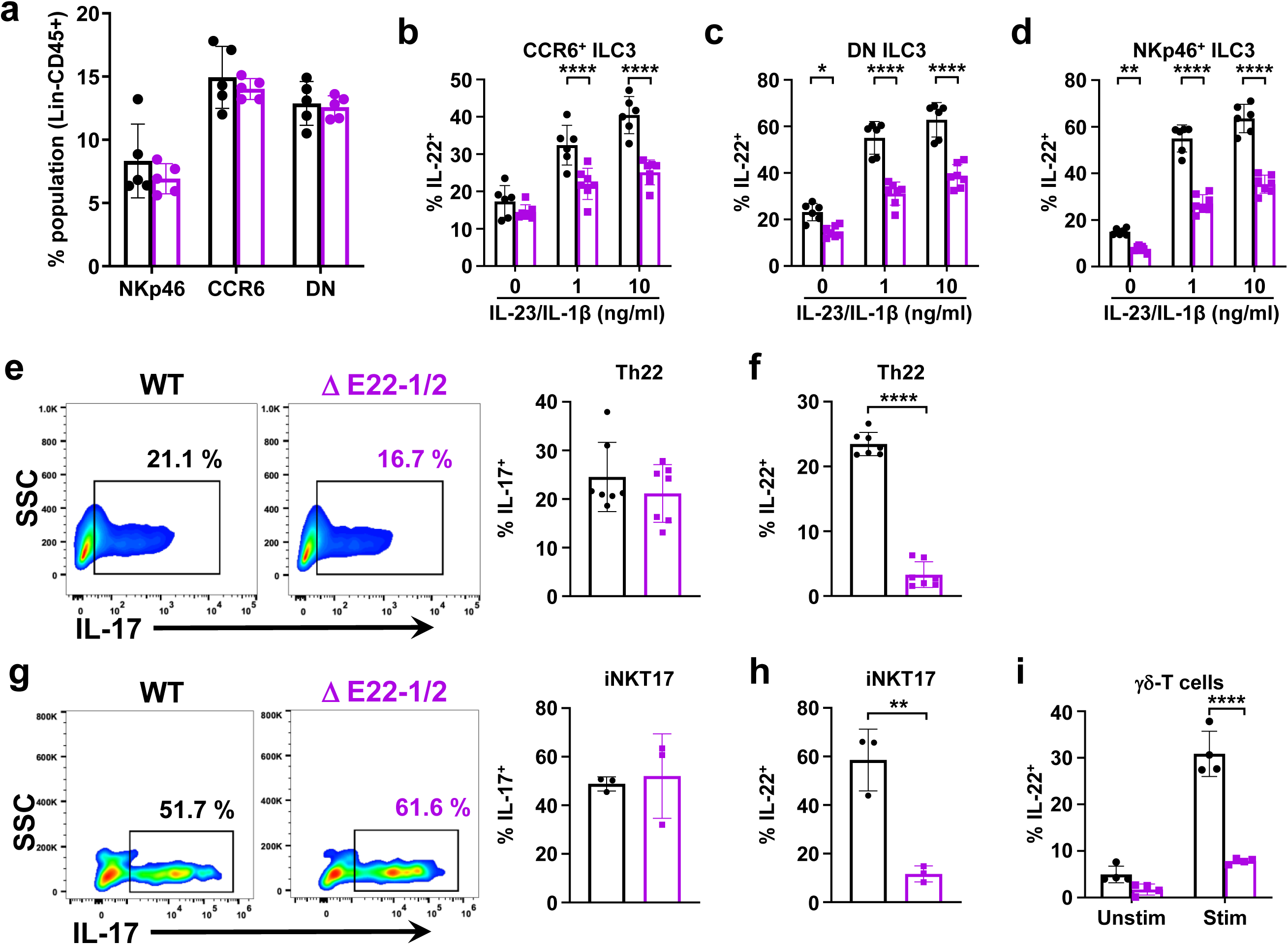
E22-1 and E22-2 regulate IL-22 expression in type 3 cells. (a) Percentage of SI-LP ILC3 subsets in WT (black) and E22-1/2^Δ^ mice (purple). (b-d) Percentages of IL-22^+^ ILC3s in all three subsets of SI-LP ILCs in WT and E22-1/2^Δ^ mice, after 3.5 h of ex vivo simulation with indicated concentrations of IL-1β+IL-23. (e) Representative flow plots (left) and percentage of IL-17^+^ (right) and of IL-22^+^ ex vivo polarized Th22 cells (f). (g) Representative flow plots (left) and percentage of IL-17^+^ (right) iNKT cells and of IL-22^+^ (h) iNKT cells isolated from lymph nodes. (i) Percentage of IL-22^+^ γδ-T cells isolated from SI-LP. Data are shown as mean ± SD and are pooled (a-f, i), or are representative (g, h) of at least two independent experiments. \**P* < 0.05; \*\**P* < 0.01; \*\*\**P* < 0.001; \*\*\*\**P* < 0.0001; two-way ANOVA with Sidak’s multiple comparisons test (a-d, i) or unpaired t-test (e-h).

**Supplemental Fig. 4:**
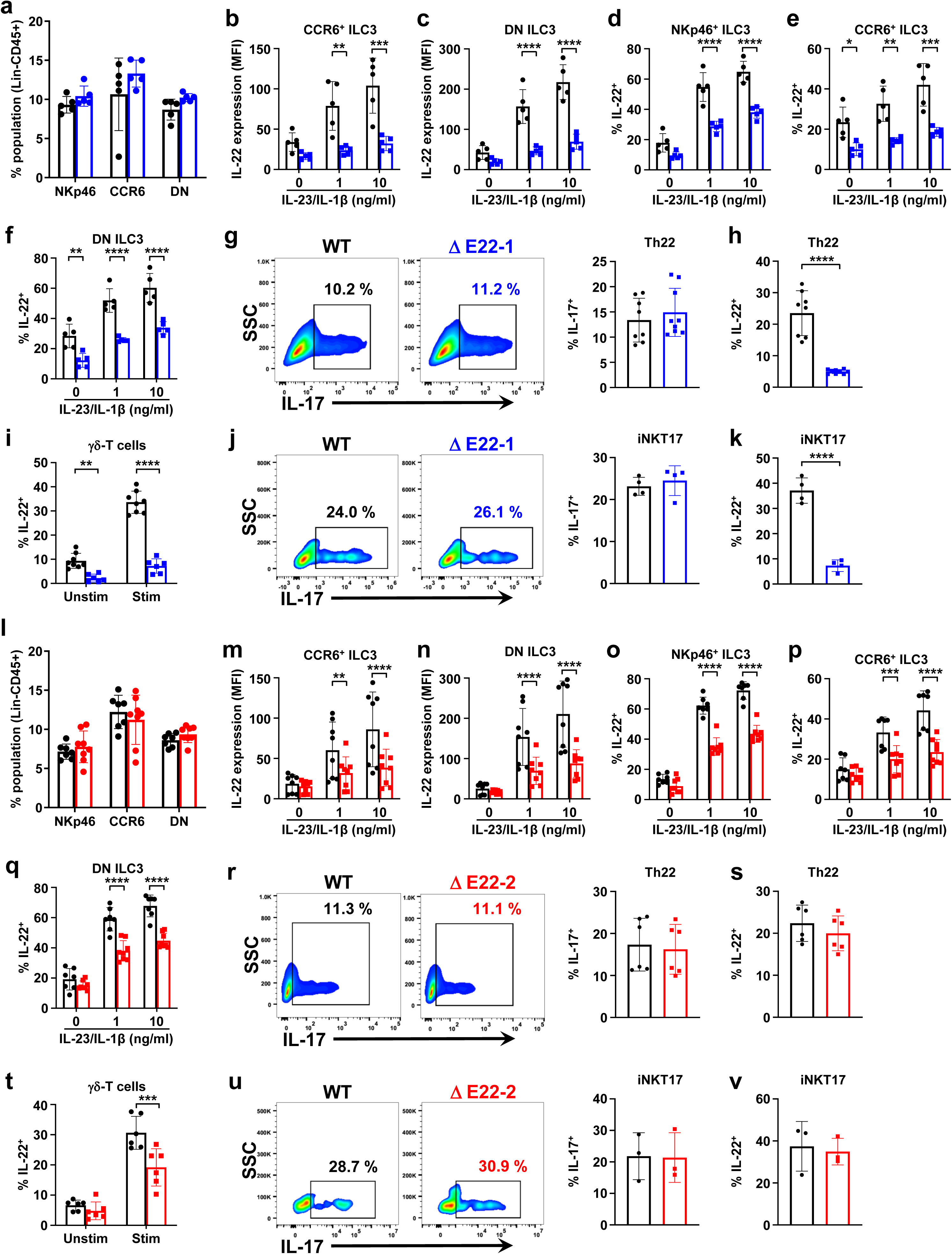
Cell type specificity of E22-1 and E22-2 cis-elements. (a,l) Frequency of NKp46^+^, CCR6^+^, and DN ILC3 subsets in SI-LP isolates. (b-f, m-q) Flow cytometry analysis of IL-22 MFI (b, c, m, n) or percentages of IL-22^+^ cells (d-f, o-q) in SI-LP ILC3 subsets treated ex vivo with indicated concentrations of IL-1β+IL-23. Representative flow plots (left) and percentages of IL-17^+^ ex vivo polarized Th22 (g, r) and NKT17 (j, u) cells. Percentage of IL-22^+^ ex vivo polarized Th22 (h, s), γδ-T (i, t), NKT17 (k, v) cells. Black - WT mice, blue - E22-1^Δ^ mice, and red - E22-2^Δ^ mice. Data are shown as mean ± SD and are pooled (a-i, l-t), or are representative (j, k, u, v) of at least two independent experiments. \**P* < 0.05; \*\**P* < 0.01; \*\*\**P* < 0.001; \*\*\*\**P* < 0.0001; two-way ANOVA with Sidak’s multiple comparisons test (a-f, i, l-q, t) or unpaired t-test (g, h, j, k, r, s, u, v).

**Supplemental Fig. 5:**
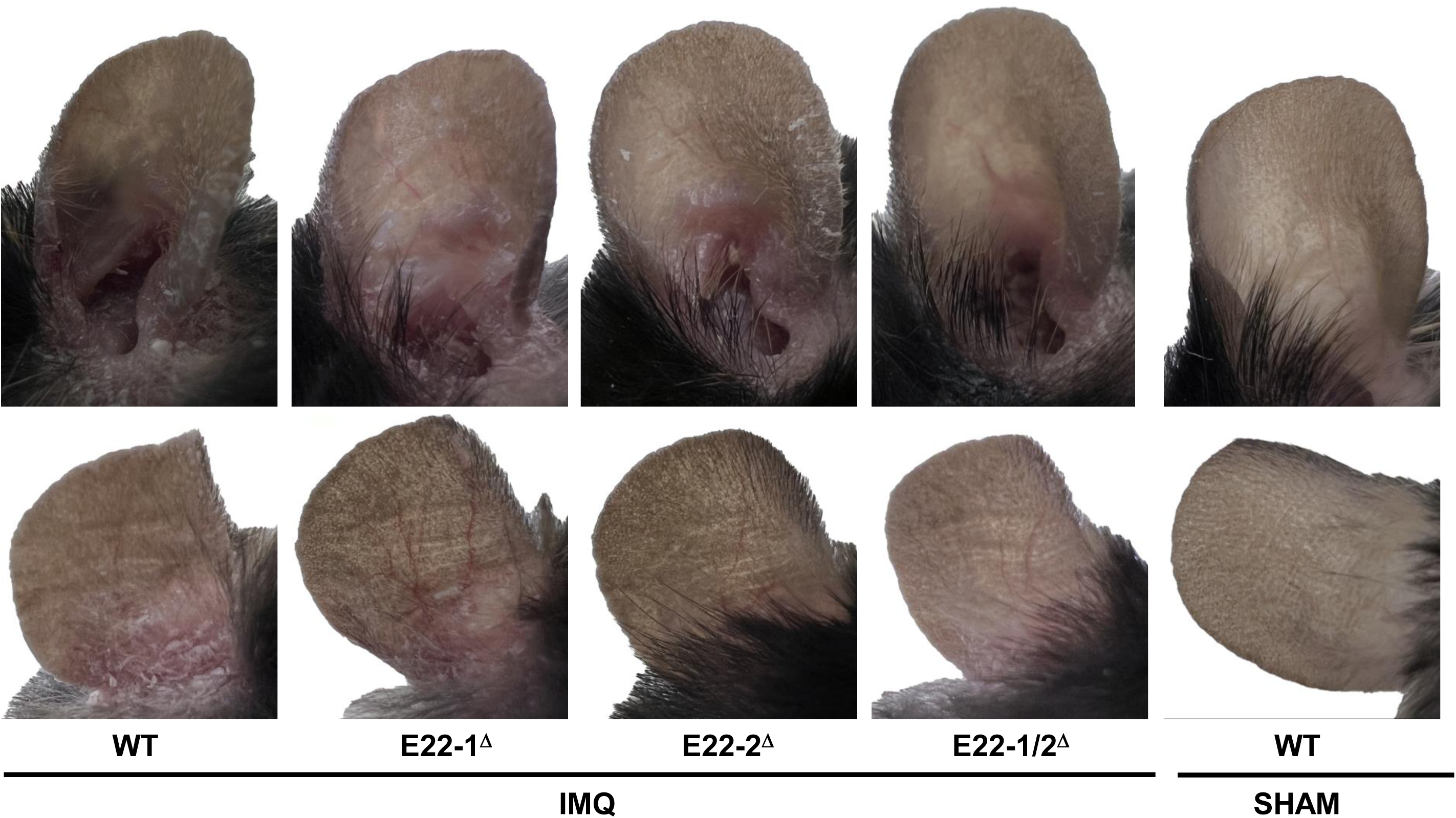
E22-1 and E22-2 driven IL-22 expression is required for the development of severe IMQ-induced psoriasis. Phenotypic presentation of mouse ear skin following 9 days of treatment with IMQ or control cream. Data are representative of three independent experiments.

**Supplemental Fig. 6:**
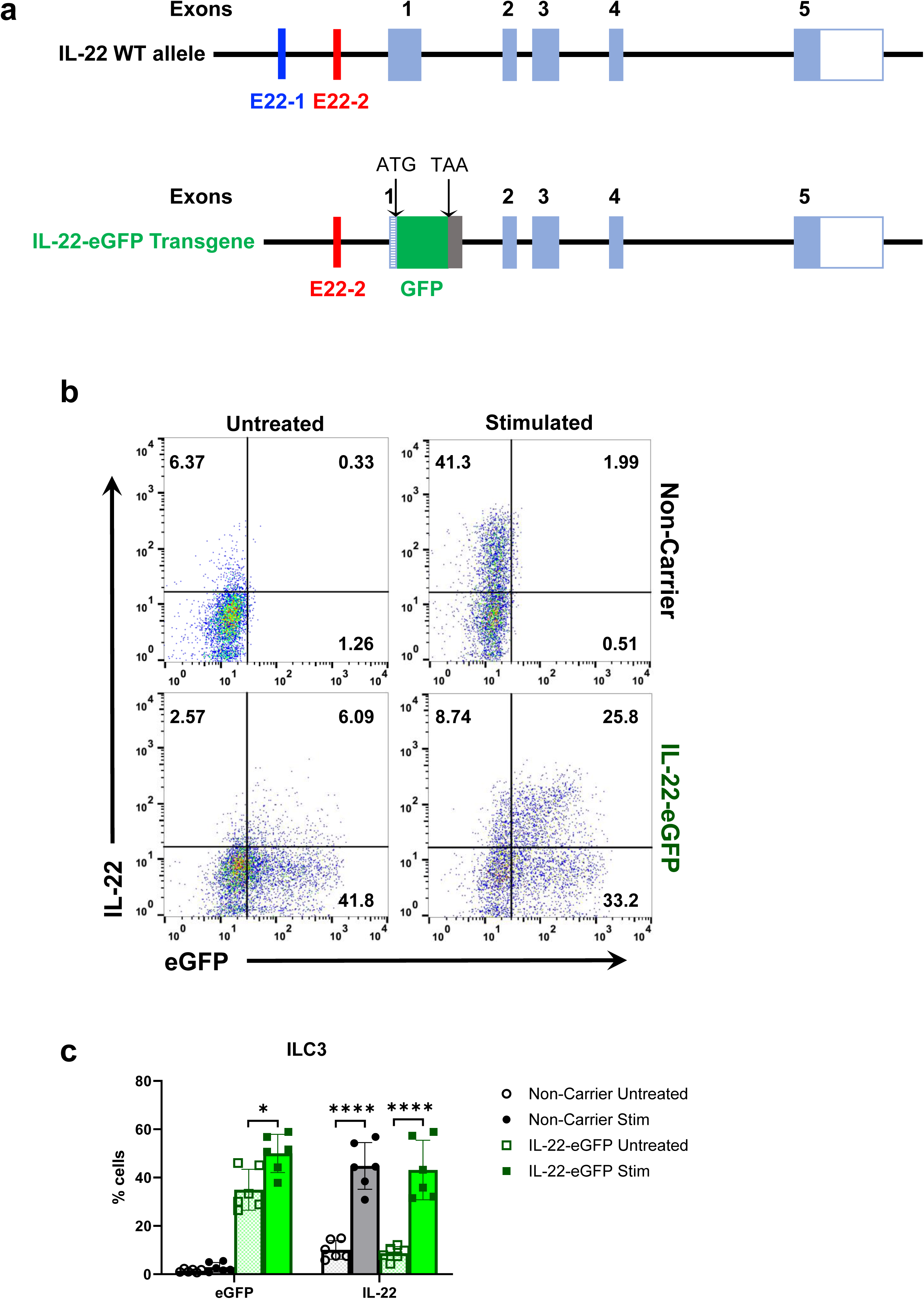
E22-2 drives IL-22 expression in ILC3s. (a) Graphic representation of the IL-22 WT allele and IL-22-eGFP transgene derived from engineering a C57BL/6 BAC clone^33^. (b, c) Representative flow plots (b) and percentages (c) of SI-LP total ILC3s showing endogenous expression of IL-22 and/or eGFP reporter in IL-22-eGFP and non-carrier mice. The SI-LP isolates were either left untreated or were stimulated with 10 ng/ml of each IL-1β and IL-23. Data are shown as mean ± SD and are pooled from two independent experiments. \**P* < 0.05; \*\*\*\**P* < 0.0001; three-way ANOVA with Tukey’s multiple comparisons test (c).

**Supplemental Fig. 7:**
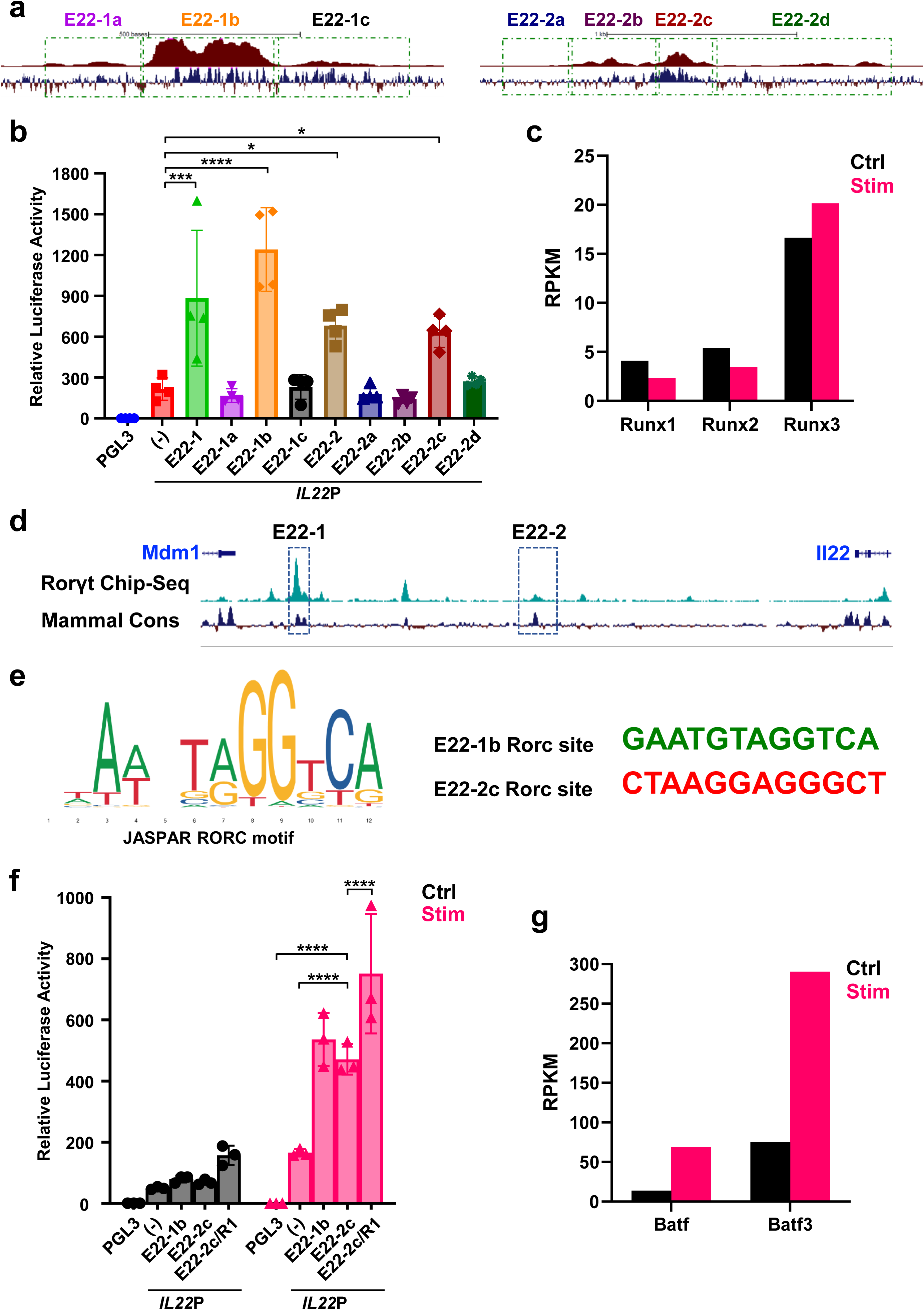
E22-1b and E22-2c are the core enhancer regions of E22-1 and E22-2, respectively. (a) Schematic showing different segments of E22-1 and E22-2 enhancers. ATAC-seq tracks of Mnk3 cells stimulated with 10 ng/ml of IL-1β and IL-23 for 16 h are displayed in maroon, and mammalian conservation in black. (b) Luciferase reporter activity of indicated segments of E22-1 and E22-2 enhancers. Mnk3 cells were nucleofected, 24 h later treated with 10 ng/ml of IL-1β and IL-23 or left untreated for 16 h. The reporter activity is relative to PGL3-Basic vector. (c) RNA-seq data analysis showing RPKM values of *Runx* factors in Mnk3 cells stimulated with 10 ng/ml of IL-1β and IL-23 or left untreated for 16 h. (d) UCSC genome browser view displaying RORγt Chip-seq^40^ track (green) in mouse Th17 cells at *Il22* locus. E22-1 and E22-2 are highlighted with blue boxes. (e) Comparison of JASPAR consensus RORγt motif with predicted sites in E22-1b and E22-2c. (f) Luciferase reporter assay comparing activity of E22-2c and with its RORγt motif replaced by the one from E22-1b (E22-2c/R1). Nuceleofected Mnk3 cells were treated 24 h post-nucleofection with 10 ng/ml of IL-1β and IL-23 or left untreated for 16 h. (g) RNA-seq data analysis showing RPKM values of *Batf* factors in Mnk3 cells stimulated for 16 h with 10 ng/ml of IL-1β and IL-23 or left untreated. Data are shown as mean ± SD and are pooled from four (b) or three (f) independent experiments. \**P* < 0.05; \*\*\**P* < 0.001; \*\*\*\**P* < 0.0001; one-way ANOVA or two-way ANOVA followed with Dunnett’s multiple comparisons test.

## Notes

### Competing Interest Statement

The authors have declared no competing interest.

